# RPRM negatively regulates ATM levels involving its phosphorylation mediated by CDK4/CDK6

**DOI:** 10.1101/2021.11.10.468148

**Authors:** Yarui Zhang, Guomin Ou, Zhujing Ye, Zhou Zhou, Qianlin Cao, Mengting Li, Jingdong Wang, Jianping Cao, Hongying Yang

## Abstract

Sensitizing cancer cells to radio- and chemotherapy remains a hot topic in cancer treatment. Here it is identified that Protein Reprimo (RPRM) negatively regulates the levels of ataxia-telangiectasia mutated (ATM) protein kinase, a master regulator of DNA damage response (DDR) in the presence of DNA double-strand breaks (DSBs), resulting in impaired DNA repair efficiency and enhanced cellular sensitivity to genotoxic agents. Mechanistically, although RPRM is primarily located in cytoplasm, it rapidly translocates to nucleus shortly after induced by X-irradiation, interacts with ATM and promotes the nuclear export and proteasomal degradation of ATM. The nuclear translocation of RPRM is associated with its phosphorylation at serine 98, which is mediated by cyclin-dependent kinases 4/6 (CDK4/6). Inhibition of CDK4/6 stabilizes RPRM and promotes its nuclear import, in turn enhances the nuclear export of ATM and the reduction of ATM levels. As a result, RPRM overexpression and its phosphorylation inhibition sensitize cells to genotoxic agents. Moreover, RPRM deficiency significantly increases resistance to radiation-induced damage both in vitro and in vivo. These findings establish a crucial regulatory mechanism in which ATM is negatively modulated by RPRM, suggesting that RPRM may serve as a novel target for both cancer therapy and radiation protection.

## Introduction

Genotoxic agents such as ionizing radiation (IR) and DNA-damaging agents are widely used in the radiotherapy and chemotherapy for cancer patients. DNA double-strand breaks (DSBs) caused by genotoxic agents are a vital determining factor for the efficacy and side effects of therapy. The ataxia-telangiectasia mutated (ATM) protein kinase that is named after Ataxia Telangiectasia (AT), an autosomal, recessive disease caused by mutations in the atm gene,^1^ is a master regulator of DNA damage response (DDR) in the presence of DSBs, which determines DNA repair efficiency and cellular sensitivity to DSBs, thus playing critical roles in genomic stability, cell death, survival, etc.^2–5^ Mechanistically, upon DSB induction in mammalian cells, the intermolecular autophosphorylation and monomerization of ATM occur rapidly, which is essential for its function in DDR.^6, 7^ ATM is also recruited to DSB sites and activated by MRE11-RAD50-NBS1 (MRN) complex.^8^ Activated ATM then phosphorylates its thousands of downstream DDR-related proteins such as H2AX, 53BP1, P53, CHK2, etc., thus setting an intricate and fine-tuned DDR network to repair DNA.^4, 5, 9, 10^ Therefore, ATM deficiency leads to enhanced cellular sensitivity to genotoxic agents. A typical example is A-T patients who exhibit hypersensitivity to IR in addition to a wide range of symptoms including neurodegeneration, senescence, immunodeficiency and cancer susceptibility.^11–14^ Due to the crucial roles it plays in DNA repair and cell killing in response to DNA damage, ATM is considered as an effective and sought-after target for cancer therapy. It has been found that KU-55933, the most widely tested ATM kinase inhibitor, increases the radiosensitivity of DAB2IP-deficient bladder cancer cells,^15^ and AZ32, a novel ATM inhibitor, can promote apoptosis in glioblastoma mouse model in combination with radiotherapy.^16^ Furthermore, its two highly selective inhibitors, AZD0156^17^ and AZD1390,^18^ have entered phase I cancer clinical trials.^19^ There have been extensive studies on how ATM regulates its various downstream targets of DDR and the underlying mechanisms,^4, 5, 9, 10^ but how ATM itself is regulated immediately after DSB induction is still not fully understood.

Reprimo (RPRM), a single-exon and intronless 327-bp gene that is located at 2q23, encodes a highly glycosylated protein.^20^ RPRM is believed to be a tumor suppressor gene.^21^ Its downregulation due to the aberrant methylation of RPRM promoter has been found in a variety of cancers such as breast cancer,^22^ gastric cancer,^23^ pituitary cancer,^24^ pancreatic cancer,^25^ etc. In addition, overexpressing RPRM inhibits the cell proliferation, colony formation, migration and invasiveness of cancer cells.^22–24, 26, 27^ RPRM expression is induced in response to DNA damage caused by X-irradiation and other genotoxic agents.^20, 23^ And upon induction, RPRM can trigger DNA damage-induced p53-dependent G2 arrest by inhibiting the activation of Cyclin B1·Cdc2 complex.^20, 28^ Furthermore, RPRM overexpression enhances apoptosis induced by DNA damage agents.^23^ All these results suggest that RPRM may be involved in DDR pathway. However, whether RPRM is integrated in ATM signaling pathway and what role it plays in DNA repair remain unknown.

In this study, we identified RPRM as a novel negative regulator of ATM. Shortly after generation of DSBs, the induced RPRM promotes the nuclear export and degradation of ATM through translocating from cytoplasm to nucleus and interacting with ATM. Moreover, cyclin-dependent kinase 4/6 (CDK4/6) are involved in the down-regulation of ATM by RPRM via its modulation on RPRM phosphorylation. Thus, we discovered the crucial functions of RPRM in DNA repair and cellular sensitivity to DNA damage, suggesting that RPRM may be a novel potential target for both cancer therapy and radiation protection.

## Results

### RPRM binds to ATM and down-regulates ATM levels

To determine the exact roles RPRM may play in DDR pathway, we firstly established a variety of cell lines with RPRM overexpression such as H460-RPRM, A549-RPRM, AGS-RPRM, SNU-1-RPRM and HaCaT-RPRM, as well as RPRM-deficient H460 and WS1 cells, pairing with their relevant negative control cells (NC) (Supplementary Fig. S1). We also established a strain of RPRM knockout mice using CRISPR-Cas9 technique, and obtained RPRM knockout mouse embryonic fibroblasts (RPRM^-/-^ MEFs). We confirmed an obvious RPRM induction after irradiation in RPRM^+/+^ MEFs (Supplementary Fig. S2a), which agreed with what was previously reported.^20^ The rapid RPRM induction by IR was also observed in different cancer cell lines with RPRM overexpression including non-small cell lung cancer cell lines and gastric cancer cell lines (Supplementary Fig. S2b, c).

Unexpectedly, by using co-immunoprecipitation assay, we found that RPRM interacted with ATM in H460-RPRM cells, and the interaction was enhanced after X-irradiation (Fig. 1a, b). Using immunofluorescent microscopy, the co-localization of RPRM and ATM was observed in irradiated RPRM^-/-^ MEFs re-expressing RPRM (Fig. 1c). But this co-localization was not seen in irradiated RPRM^-/-^ MEFs transfected with NC vectors (Supplementary Fig. S3). Moreover, compared with H460-NC cells, H460-RPRM cells exhibited decreased levels of the total ATM protein, this reduction was especially significant at 2 h after X-irradiation and last for at least another 4 hours, accompanied by a reduction in phosphorylated ATM (Fig. 1d, e). Similar results were also observed in HaCaT human skin keratinocytes (Fig. 1f), A549 lung cancer cells, AGS and SNU-1 gastric cancer cells (Supplementary Fig. S4). These data suggest that RPRM down-regulates ATM levels after radiation exposure. To further confirm this, we also used RPRM^-/-^ MEFs to investigate how ATM levels would be changed by RPRM, and found that ATM levels were generally higher in RPRM^-/-^ MEFs than in WT MEFs before and after irradiation (Fig. 1g, h), implying that RPRM knockout increased ATM levels. However, this trend was reversed when RPRM was re-expressed in RPRM^-/-^ MEFs and exposed to IR (Fig. 1i, j). All these data indicate that X-induced RPRM negatively regulates ATM levels.

**Figure 1.**
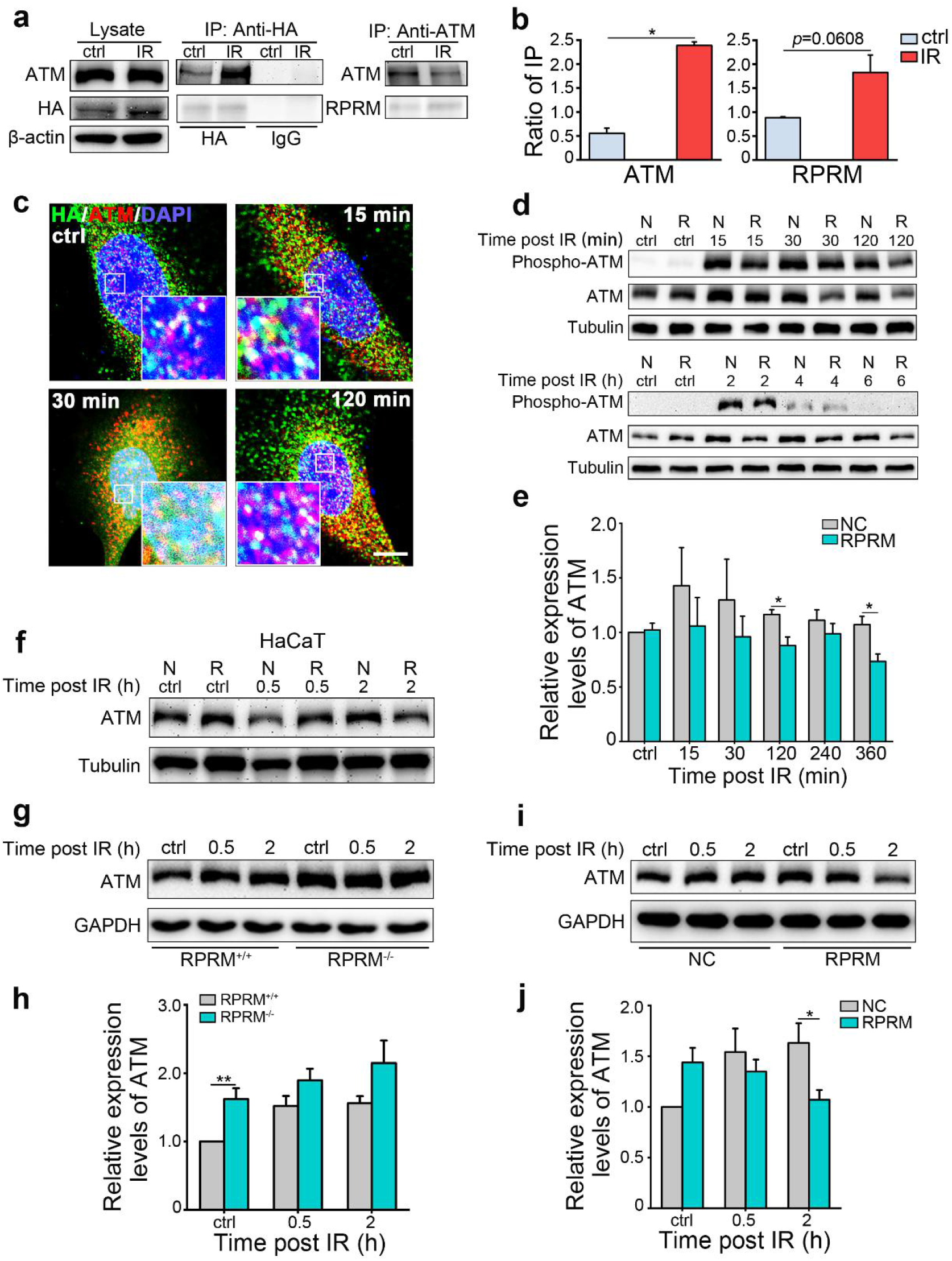
RPRM binds to ATM and down-regulates ATM levels. **a, b** Co-immunoprecipitation assay (Co-IP) was performed to demonstrate the interaction between ATM and RPRM, which was enhanced in irradiated H460-RPRM cells. **c** Representative immunofluorescence images of co-localization of RPRM and ATM in RPRM^-/-^ MEFs reexpressing RPRM at different times post 20 Gy X-irradiation. **d, e** Change in the levels of total ATM and phospho-ATM in H460-NC (N)/RPRM (R) cells at different times post 2 Gy X-irradiation. Quantification of ATM levels (**e**), which were the average of three independent experiments. **f** Change in the ATM levels of HaCaT-NC (N)/RPRM (R) cells irradiated with 2 Gy X-rays showed that RPRM overexpression in HaCaT cells decreased ATM levels upon IR. **g, h** The ATM levels of RPRM^+/+^ MEFs were lower than those of RPRM^-/-^ MEFs at different times post 20 Gy X-irradiation. Bar graph (**h**) shows quantification of ATM levels. **i, j** RPRM reexpression reduced the ATM levels in RPRM^-/-^ MEFs post 20 Gy X-irradiation. Bar graph (**j**) shows quantification of ATM levels. Experiments were independently repeated three times with similar results. \**P* < 0.05; \*\**P* < 0.01.

### RPRM promotes ATM nuclear export and degradation

We further investigated how RPRM down-regulated ATM. We did not observe any significant reduction in ATM expression in irradiated cells with RPRM overexpression at mRNA level (Supplementary Fig. S5a), suggesting that RPRM does not regulate ATM at transcription level. At protein level, in the presence of cycloheximide, a protein synthesis inhibitor, the ATM protein levels in H460-RPRM cells were significantly lower than those in H460-NC cells regardless of radiation exposure. The difference between H460-RPRM and H460-NC cells was greater after irradiation (Fig. 2a, b; Supplementary Fig. S5b). The same result was observed in AGS-RPRM cells (Supplementary Fig. S5c). But this reduction of ATM levels in RPRM-overexpressed cells compared with NC cells was no longer observed when the cells were treated with MG132, a proteasome inhibitor, prior to irradiation (Fig. 2c, d). These data suggest that RPRM down-regulates ATM at protein level through promoting its degradation but not through inhibiting its synthesis. Moreover, we used RPRM^-/-^ MEFs to further confirm the effect of RPRM on ATM degradation. As shown in Figure 2e and f, the ATM levels of RPRM^-/-^ MEFs were slightly higher than those of WT MEFs with and without radiation exposure when the protein synthesis was inhibited, although the difference did not reach statistical significance. However, re-expression of RPRM in RPRM^-/-^ MEFs decreased the ATM levels, and the reduction was more significant after irradiation (Fig. 2g, h), suggesting that RPRM promotes ATM degradation, resulting in a reduction of ATM levels, and this promotion is more obvious after irradiation.

**Figure 2.**
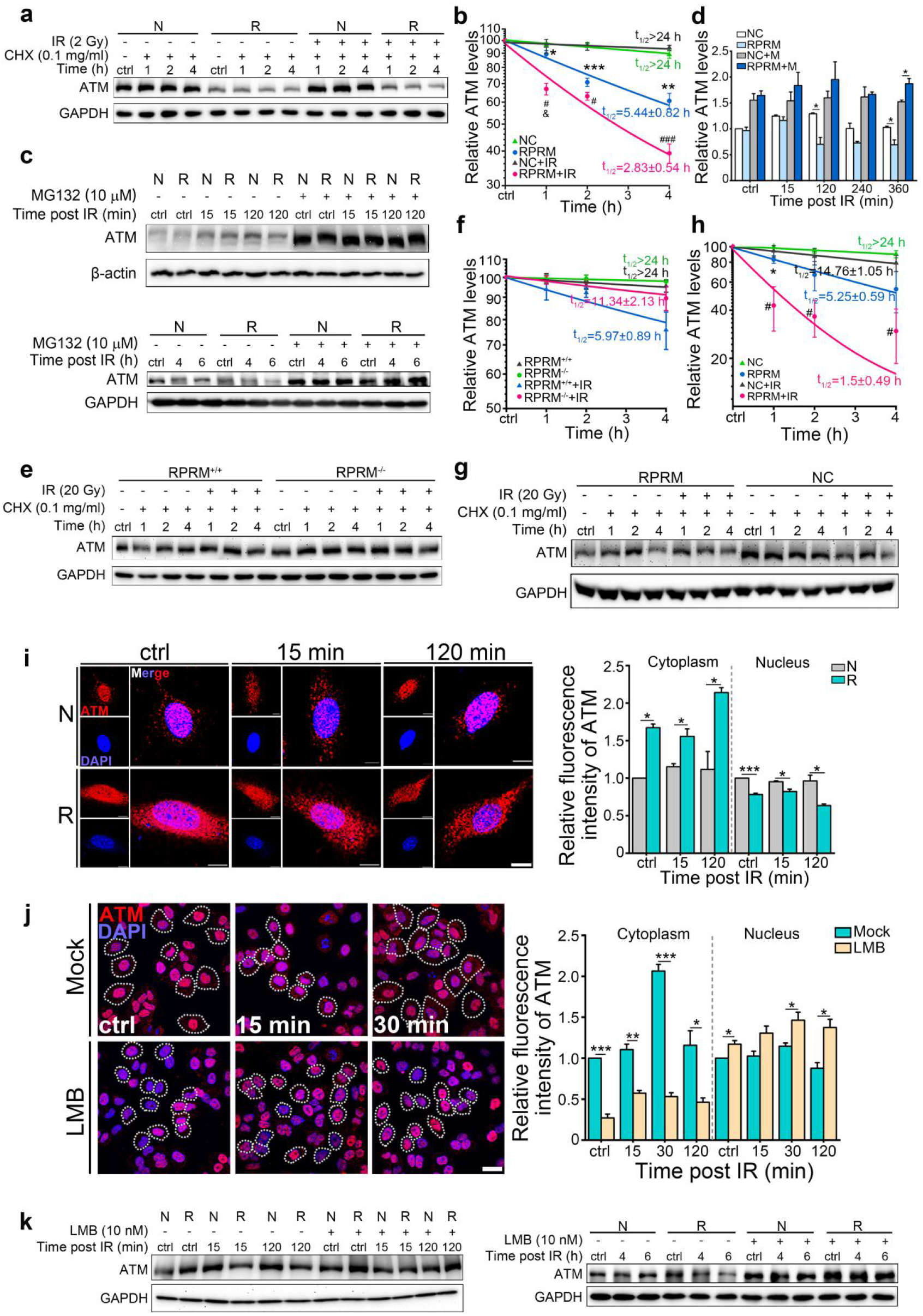
RPRM promotes the nuclear export and degradation of ATM. **a, b** Kinetics of the change in the ATM levels of H460-NC (N)/RPRM (R) cells with and without irradiation in the presence of cycloheximide (CHX, 0.1 mg/ml) examined by immunoblotting. (*, RPRM vs NC; #, RPRM+IR vs NC+IR; &, RPRM+IR vs RPRM). **c, d** Change in the levels of ATM of H460-NC (N)/RPRM (R) cells at the indicated time points after 2 Gy X-irradiation with and without MG132 (10 μM) pre-treatment. **e-h** Kinetics of the change in ATM levels of RPRM^+/+^ MEFs, RPRM^-/-^ MEFs (**e, f**), RPRM^-/-^ MEFs re-expression NC/RPRM (**g, h**) with and without irradiation in the presence of cycloheximide (0.1 mg/ml) examined by immunoblotting. (*, RPRM vs NC; #, RPRM+IR vs NC+IR). **i** Change in the subcellular localization of ATM in RPRM^-/-^ MEFs re-expressing NC (N) and RPRM (R) after 20 Gy X-irradiation. Scale bars, 10μm. **j** Representative immunofluorescence images of ATM in H460-RPRM cells pretreated with leptomycin B (LMB, 10 nM) for 1 h at different times post 2 Gy X-irradiation. White dotted circles indicate cytoplasmic ATM. Quantification of the fluorescence intensity of the nuclear and cytoplasmic ATM. Scale bar, 20 μm. **k** The effect of leptomycin B (LMB, 10 nM) pretreatment on the total ATM levels of H460-NC/RPRM cells at different times post 2 Gy X-irradiation. Experiments were independently repeated three times with similar results. \**P* < 0.05; \*\**P* < 0.01; \*\*\**P* < 0.001; #*P* < 0.05; ###*P* < 0.001; &*P* < 0.05.

Most proteins can be degraded through ubiquitin-proteasome pathway, which may occur in both cytoplasm and nucleus.^29^ Although ATM is well known as a nuclear protein in dividing cells, missense mutations can lead to its abnormal cytoplasmic localization, which is correlated with its decreased expression.^30^ Thus, we hypothesized that RPRM could enhance the nuclear export of ATM after exposed to IR, in turn promoting its degradation and leading to its down-regulation. By using immunofluorescent microscopy, we clearly observed the nucleus-to-cytoplasm translocation of ATM in irradiated NC cells. Most importantly, RPRM overexpression obviously increased the levels of cytoplasmic ATM, especially after radiation, accompanied with decreased levels of nuclear ATM (Supplementary Fig. S6a). But interestingly, the phospho-ATM still predominantly accumulated in nucleus (Supplementary Fig. S6b). The ATM translocation was also confirmed by immunoblotting on the cytoplasmic and nuclear fractions of cells, which clearly showed an increase of the cytoplasmic ATM levels and a reduction of the nuclear ATM levels in RPRM-overexpressed cells after IR (Supplementary Fig. S6c). In contrast, no obvious nuclear-cytoplasmic translocation of ATM was observed in irradiated cells when RPRM was knocked out. And re-expression of RPRM significantly increased the cytoplasmic ATM levels and reduced the nuclear ATM levels before and after IR (Fig. 2i). Moreover, when leptomycin B (LMB), a widely used nuclear export inhibitor, was added into the culture of RPRM-overexpressed cells, the cytoplasmic ATM levels were dramatically decreased while the nuclear ATM levels were increased (Fig. 2j), which was very likely due to the nuclear export of ATM being diminished by LMB. At the same time, no reduction of the total ATM levels was detected (Fig. 2k; Supplementary Fig. S6d). These data indicate that RPRM enhances the nuclear export of ATM after IR, thus resulting in the reduction of the nuclear and total ATM levels.

### RPRM translocates from cytoplasm to nucleus upon irradiation, which involves CDK4/6

RPRM has been thought to be primarily located in cytoplasm since it was first reported.^20^ However, our discovery that RPRM interacted with ATM and promoted its nuclear export implied that RPRM may translocate from cytoplasm to nucleus due to the major nuclear localization of ATM. We then explored whether IR would change the subcellular localization of RPRM. Interestingly, cells with RPRM overexpression showed a significant increase in the immunofluorescence staining of RPRM in nucleus 15 and 30 min after IR, but this increase was not observed by 2 h post IR (Fig. 3a), indicating an occurrence of quick nuclear translocation of RPRM after its induction by IR. This nuclear translocation of RPRM was also confirmed by western blotting on the nuclear fractions of cells (Fig. 3b). These data demonstrate that RPRM translocates from cytoplasm to nucleus shortly after irradiation.

**Figure 3.**
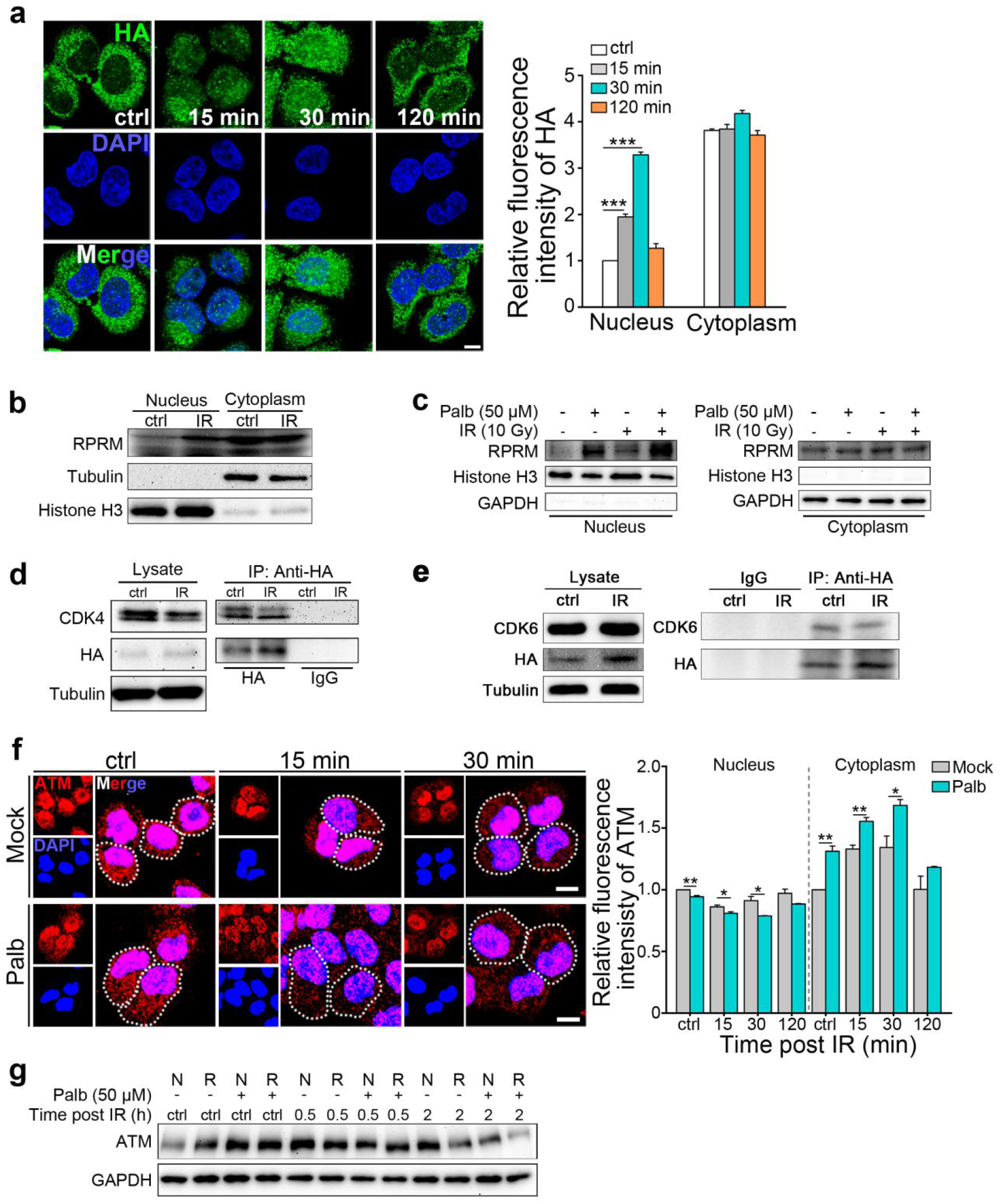
RPRM translocates from cytoplasm to nucleus upon irradiation, which involves CDK4/6. **a** Representative immunofluorescence images showing RPRM translocation from cytoplasm to nucleus in H460-RPRM cells shortly after 10 Gy X-irradiation and quantification of the fluorescence intensity of the nuclear and cytoplasmic RPRM. Scale bar, 10 μm. **b** Immunoblotting on the nuclear and cytoplasmic RPRM of H460-RPRM cells 30 min after 10 Gy X-irradiation confirmed the nuclear translocation of RPRM upon IR. **c** Immunoblotting on the nuclear and cytoplasmic RPRM confirmed that the nuclear translocation of RPRM was enhanced by Palbociclib (Palb, 50 μM) pretreatment. **d, e** Co-IP result showing an interaction between RPRM and CDK4/6 in H460-RPRM cells. **f** Inhibition CDK4/6 promoted ATM nuclear export regulated by RPRM. Representative immunofluorescence images showing the effect of CDK4/6 inhibition by Palbociclib (Palb, 50 μM) pretreatment on the ATM cytoplasmic translocation in H460-RPRM cells upon 2 Gy X-irradiation and quantification of the fluorescence intensity of the nuclear and cytoplasmic ATM. Scale bar, 10 μm. **g** Immunoblotting results confirmed that CDK4/6 inhibitor Palbociclib reduced ATM levels in H460-NC/RPRM cells upon 2 Gy X-irradiation. Experiments were independently repeated three times with similar results. \**P* < 0.05; \*\**P* < 0.01; \*\*\**P* < 0.001.

We further investigated how the nuclear translocation of RPRM was mediated. Interestingly, pretreating cells with Palbociclib, a highly specific inhibitor of Cyclin-dependent kinase 4/6 (CDK4/6), prior to IR enhanced RPRM nuclear translocation, even Palbociclib treatment alone could significantly increase the nuclear import of RPRM (Fig. 3c; Supplementary Fig. S7a), suggesting that CDK4/6 may play an important role in RPRM nuclear translocation. CDK4 has been found to interact with RPRM in a large-scale protein-protein interaction mapping by mass spectrometry.^31^ Using co-immunoprecipitation assay, we confirmed that CDK4 indeed bound to RPRM in H460-RPRM cells and the interaction between CDK4 and RPRM was weakened after X-irradiation (Fig. 3d). In addition, we found that CDK6 also interacted with RPRM and the CDK6-RPRM interaction was decreased after irradiation (Fig. 3e). This may be associated with the decreased levels of CDK4/6 in irradiated H460-RPRM cells (Supplementary Fig. S7b, c), which coincided with the nuclear translocation of RPRM (Fig. 3a). Furthermore, with Palbociclib, the cytoplasmic ATM levels of cells were significantly increased before and post IR while the nuclear ATM levels were obviously decreased (Fig. 3f). Moreover, the total ATM levels were markedly reduced by Palbociclib in both RPRM-overexpressed cells and NC cells 2 h after exposed to X-rays (Fig. 3g). This indicated that CDK4/6 inhibition promoted the ATM nuclear export and decreased the ATM levels. Taken together, these data suggest that CDK4/6 are involved in the nuclear translocation of RPRM and its negatively regulatory effect on ATM.

### CDK4/6 phosphorylates RPRM on serine 98, the unphosphorylation status of RPRM is critical for its stabilization and nuclear translocation

Since RPRM contains a predicated phosphorylation site at serine 98,^21^ and CDK4/6 are members of Ser/Thr protein kinase family, we then performed mass spectrometry analysis to determine whether CDK4/6 could directly phosphorylate RPRM. Not unexpectedly, CDK4/6 indeed phosphorylated RPRM at serine 98 (Fig. 4a, b), indicating that CDK4/6 regulates RPRM via a post-translational mechanism. Moreover, after CDK4/6 inhibition by Palbociclib, the RPRM levels of cells were obviously increased, which was similar to the change in the presence of MG132 (Fig. 4c), suggesting that unphosphorylated RPRM is <u>more stable</u>. To further explore whether the phosphorylation of RPRM at serine 98 mediated by CDK4/6 was critical to its nuclear translocation and negatively regulatory effect on ATM, we generated RPRM mutant (S98A) and deletion mutant (Δ(79-109)) constructs without the phosphorylation site (Fig. 4d), and transiently expressed HA-tagged full-length RPRM and RPRM mutant constructs in RPRM^-/-^ MEFs. Both mutants translocated from cytoplasm to nucleus shortly after IR just like full-length RPRM did (Fig. 4e). But the truncated mutant exhibited much less nuclear accumulation compared with the RPRM-S98A mutant, implying that the intricate mechanism of RPRM translocation may involve its phosphorylation modification as well as its structural integrity of C-terminal region. Interestingly, both RPRM-S98A and deletion mutants interacted with ATM (Fig. 4e, f), suggesting that C-terminal repeats (79-109) may not be essential to the interaction between ATM and RPRM. In agreement with the robust nuclear translocation of the RPRM-S98A mutant and its interaction with ATM, the ATM protein levels of H460-S98A cells decreased more significantly compared with H460-NC/RPRM cells under DNA damage stress such as IR and cisplatin treatment (Fig. 4g-i). These data suggest that CDK4/6 phosphorylate RPRM at serine 98, but preventing RPRM phosphorylation makes it more stable and more easily to translocate to nucleus, thus resulting in a more significant reduction in ATM levels (Fig. 4j).

**Figure 4.**
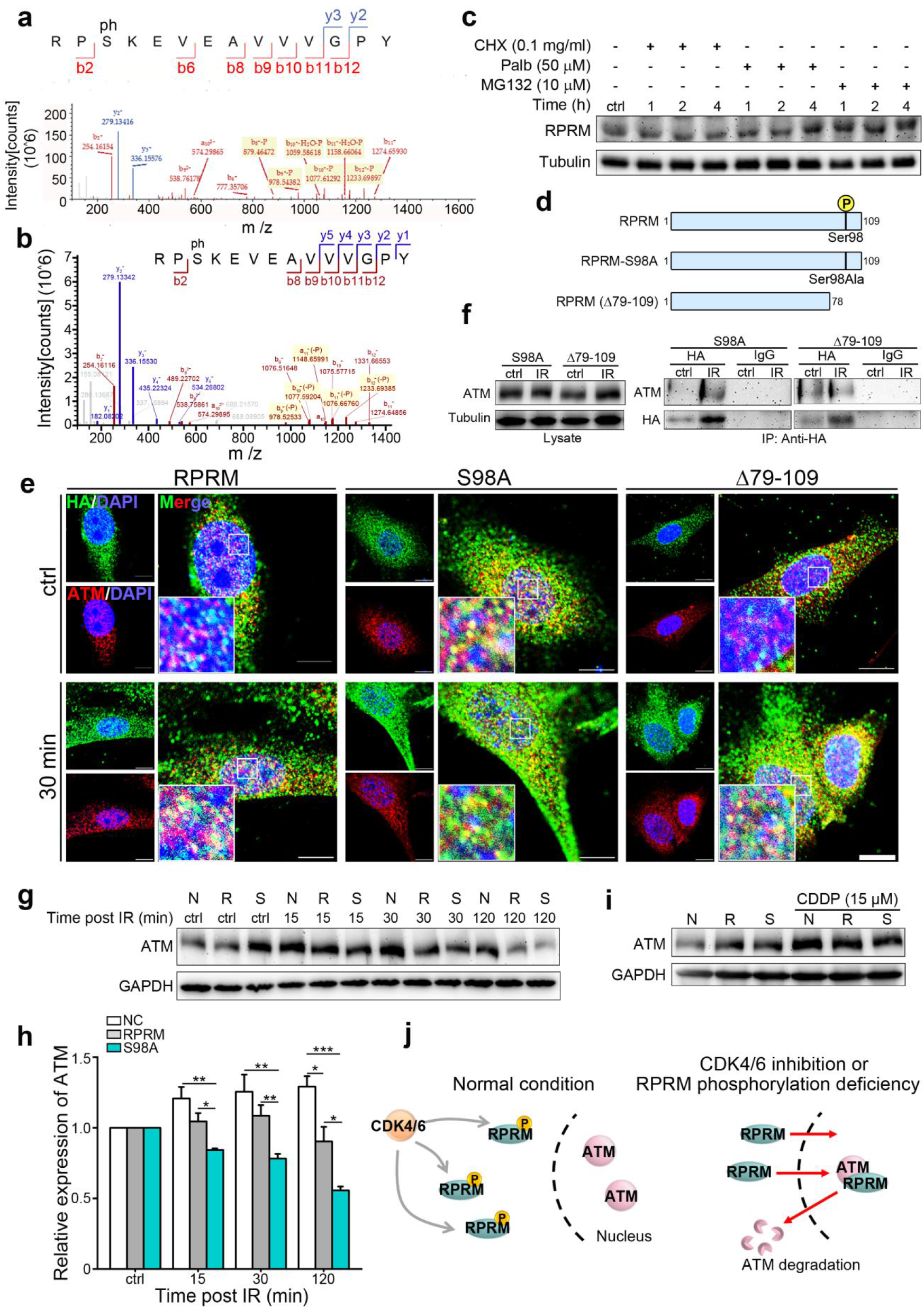
CDK4/6 phosphorylate RPRM on serine 98, but the unphosphorylation status of RPRM is critical for its stabilization and nuclear translocation. **a, b** Mass spectrometry analysis of the phosphorylation site of RPRM corresponding to CDK4 (**a**) and CDK6 (**b**). **c** Time-course analysis of the RPRM levels of H460-RPRM cells pretreated with different inhibitors. **d** Diagrams of full-length RPRM protein and the corresponding mutants. **e** Representative immunofluorescence images of the RPRM^-/-^ MEFs exogenously expressing fulllength RPRM, RPRM-S98A and ΔRPRM(79-109) at 30 min post 20 Gy X-irradiation. Scale bars, 10 μm. **f** Co-IP assay confirmed an interaction between ATM and the RPRM-S98A and truncated mutant Δ(79-109) in H460 cells. **g, h** Change in the ATM levels of H460-NC/RPRM/S98A cells at different times after 2 Gy X-irradiation. **i** Immunoblotting on the ATM expression of H460-NC/RPRM/S998A cells with or without cisplatin (CDDP, 15 μM) treatment for 24 h. **j** Working model of the regulatory effect of RPRM on ATM involving CDK4/6. Experiments (**c**, **e**-**i**) were independently repeated three times with similar results. \**P* < 0.05; \*\**P* < 0.01; \*\*\**P* < 0.001.

### RPRM inhibits DNA repair signaling pathway and enhances cellular radiosensitivity

Due to the vital role of ATM in DNA DSB repair, to investigate the consequences of the regulatory effect of RPRM on ATM, we then examined several important DSB repair indicators such as γ-H2AX foci, a widely used DNA DSB marker,^32^ 53BP1, a critical protein for triggering non-homologous end joining (NHEJ) repair in DDR,^33^ and RAD51, a vital protein in catalyzing the core reactions of homologous recombination (HR) repair.^34^ Both H2AX and 53BP1 are downstream substrates of ATM.^35^ RAD51 foci formed in nuclei after radiation exposure represent repair foci through HR,^36^ in which ATM plays an important role.^37^ As shown in Fig. 5, there was an elevated background of all three types of DNA repair foci in H460-RPRM cells compared with H460-NC cells, suggesting an elevated level of spontaneous DNA damage and repair caused by RPRM overexpression. However, at least within 2 h after irradiation H460-RPRM cells formed less γ-H2AX foci than H460-NC cells, but by 5 h post IR there were significantly more γ-H2AX foci in H460-RPRM cells than in H460-NC cells, and this trend last up to 24 h post IR (Fig. 5a; Supplementary Fig. S8a), suggesting that there existed more DSBs in irradiated cells with RPRM overexpression at longer time after IR. 53BP1 foci in H460-RPRM cells were also less than in H460-NC cells within 5 h after irradiation, although this trend was reversed by 8 h post irradiation, suggesting defective NHEJ pathway in H460-RPRM cells (Fig. 5b; Supplementary Fig. S8b). Similarly, IR-induced RAD51 foci within 8 h after irradiation were much less in H460-RPRM cells, but by 24 h post IR, H460-RPRM cells again exhibited considerably greater number of RAD51 foci than H460-NC cells (Fig. 5c; Supplementary Fig. S8c), implying defective HR pathway in H460-RPRM cells. And the results indicating that these DSB repair-related proteins were all depressed in irradiated H460-RPRM cells were in agreement with the decreased total ATM levels and weakened ATM activation (Fig. 1d, e). To further confirm that RPRM did inhibit DNA DSB repair, we also directly examined the DNA DSB accumulation in cells at different times post IR by performing comet assay. As expected, compared with NC cells, the cells overexpressing RPRM showed more severe DNA DSB accumulation shortly after IR and more DSB residues longer time after IR (Fig. 5d, e), demonstrating that RPRM drastically attenuated DNA damage repair. All these data suggest that RPRM inhibits DNA damage repair signaling pathway probably via its down-regulatory effect on ATM (Fig. 5f).

**Figure 5.**
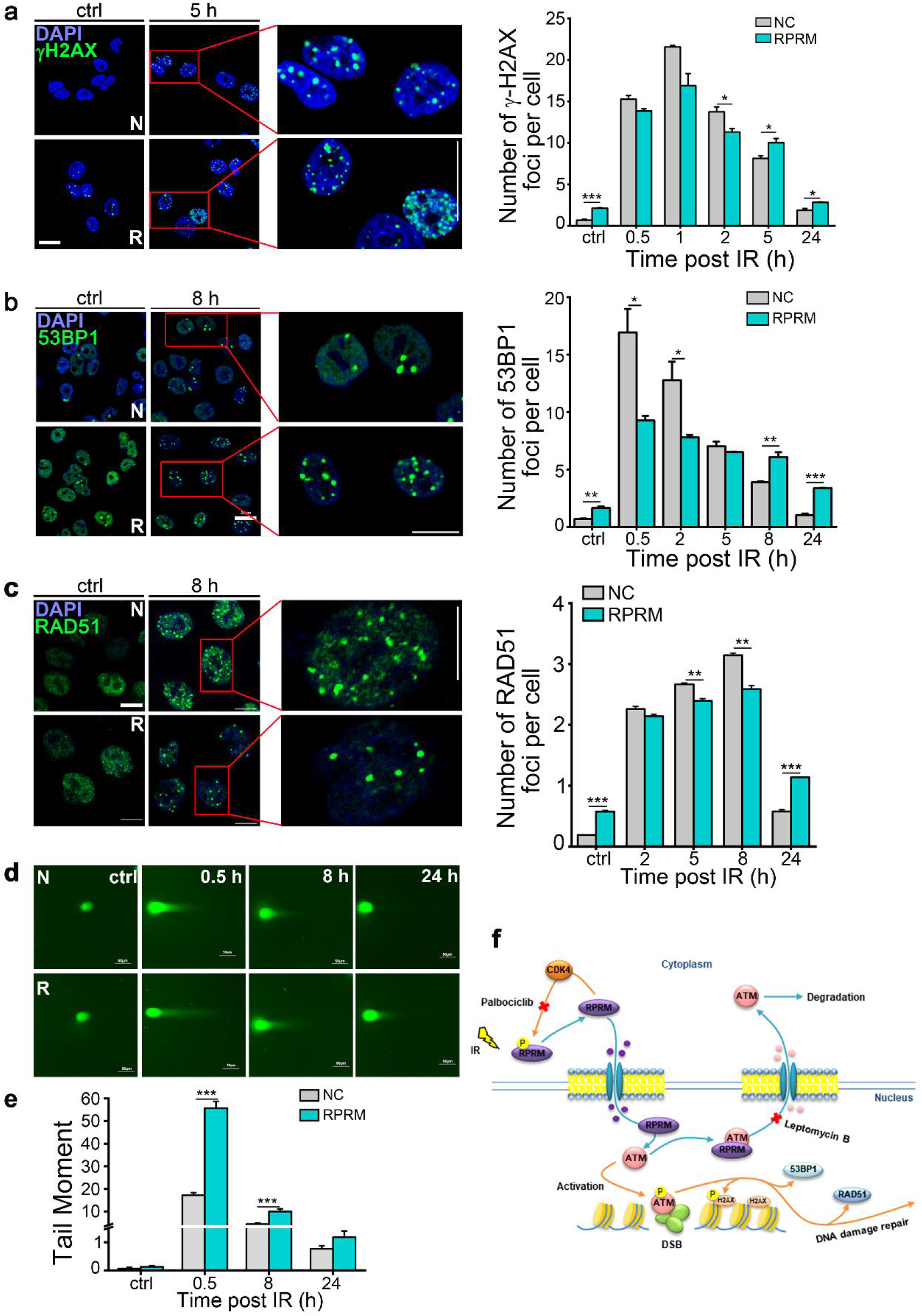
RPRM inhibits DNA repair signaling pathway. **a** Representative immunofluorescence images of γ-H2AX in H460-NC (N)/RPRM (R) cells at different times post 2 Gy X-irradiation and quantification of the number of γH2AX foci per cell. At least 200 nuclei were scored. Scale bars, 20 μm. **b** Immunofluorescence images of 53BP1 foci after cells were X-irradiated with 2 Gy and quantification. At least 200 nuclei were scored. Scale bar, 20 μm. **c** Immunofluorescence images of RAD51 foci post IR and quantification. Scale bars, 10 μm. At least 500 nuclei for each sample were scored in each experiment. **d, e** Comet assay on H460-NC/RPRM cells at different times post 2 Gy X-irradiation. The tail moments were quantified (**e**) using CometScore. At least 500 cells were scored per sample. Scale bars, 50 μm. **f** Working model of the regulatory effect of RPRM on ATM and DNA repair, leading to increased sensitivity to DNA damage. Experiments were independently repeated three times with similar results. \**P* < 0.05; \*\**P* < 0.01; \*\*\**P* < 0.001.

Furthermore, overexpression of RPRM significantly increased the radiosensitivity of cancer cells manifesting as a reduction of colony-forming ability (Fig. 6a; Supplementary Fig. S9a, b) and an increase in micronucleus formation (Fig. 6b; Supplementary Fig. S9c) upon irradiation when compared with NC cells. But LMB pre-treatment completely eliminated the increase of micronucleus formation in irradiated H460 cells caused by RPRM overexpression (Fig. 6c), suggesting that the nuclear export of ATM was essential to the radiosensitizing effect of RPRM. Conversely, RPRM knockdown significantly increased the plating efficiency of H460 cells (Fig. 6d; Supplementary Fig. S9d) and decreased the micronucleus formation of H460 and WS1 cells (Fig. 6e, f) after IR, indicating that RPRM knockdown reduced cellular radiosensitivity. Moreover, while RPRM^-/-^ MEFs formed much less micronuclei after radiation exposure than WT MEFs did (Fig. 6g), re-expression of RPRM in RPRM^-/-^ MEFs completely reversed the trend and significantly increased the micronucelus formation of RPRM^-/-^ MEFs (Fig. 6h). All these data indicate that RPRM plays a determining role in cellular radiosensitivity.

**Figure 6.**
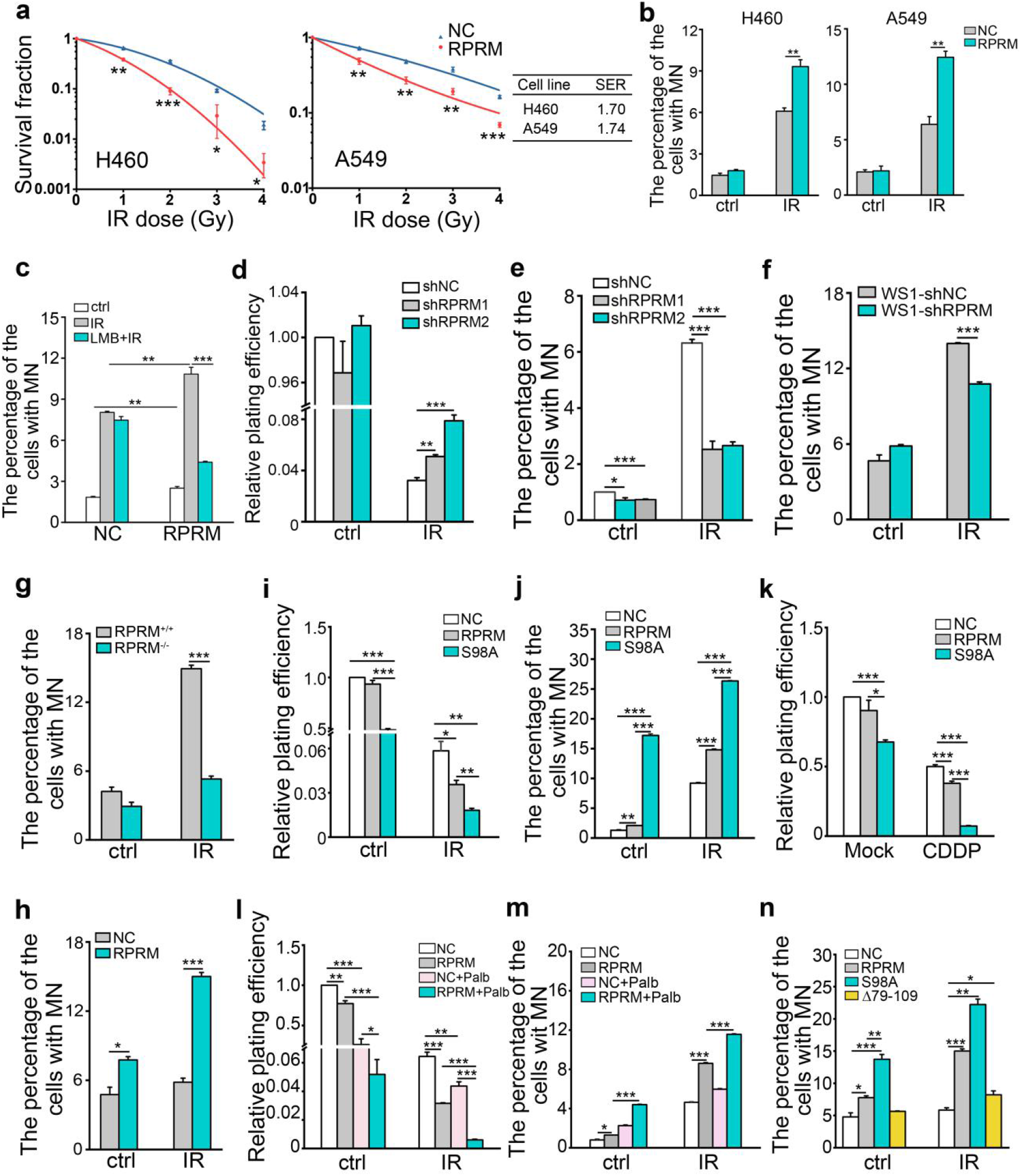
RPRM overexpression and inhibition of RPRM phosphorylation confer cellular sensitivity to DNA damage. **a** RPRM overexpression decreased the colony formation ability of cancer cells. Sensitization enhancement ratio (SER) was calculated according to the following formula: the radiation dose required for the control group to obtain 30%cell survival/the radiation dose required to obtain the same cell survival rate after overexpression of RPRM. **b** RPRM overexpression increased micronucleus formation in cancer cells upon 2 Gy X-irradiation. **c** Effect of leptomycin B (LMB, 10 nM) on the micronucleus formation of H460-NC/RPRM cells irradiated with 2 Gy X-rays. **d, e** RPRM knockdown significantly increased cellular radioresistance manifesting as increased plating efficiency (**d**) and decreased micronucleus formation (**e**) in irradiated H460 cells. Radiation doses were 4 Gy (**d**) and 2 Gy (**e**), respectively. **f** RPRM knockdown significantly reduced the micronucleus formation in WS1 cells upon 2 Gy X-irradiation. **g** RPRM knockout significantly reduced the micronucleus formation in MEFs upon 5 Gy X-irradiation.**h** Micronucleus formation of RPRM^-/-^ MEFs transfected with NC and WT RPRM plasmids after exposed to 5 Gy X-irradiation. **i** Relative plating efficiency of H460-NC/RPRM/S98A cells treated with 4 Gy X-rays. **j** Overexpression of both RPRM and RPRM-S98A increased micronucleus formation in H460 cells upon 2 Gy X-rays. **k** Relative plating efficiency of H460-NC/RPRM/S98A cells treated with 15 μM CDDP. **l** The effect of CDK4/6 inhibition by Palbociclib (Palb, 50 μM) pretreatment on the plating efficiency of H460-NC/RPRM cells upon 4 Gy X-rays. **m** The effect of CDK4/6 inhibition by Palbociclib (Palb, 50 μM) pretreatment on the micronucleus formation of H460-NC/RPRM cells after 2 Gy X-irradiation. **n** Micronucleus formation of the RPRM^-/-^ MEFs transfected with NC, full-length RPRM, RPRM-S98A and ΔRPRM(79-109) plasmids after exposed to 5 Gy X-irradiation. Experiments were independently repeated three times with similar results. \**P* < 0.05; \*\**P* < 0.01; \*\*\**P* < 0.001.

Since the phosphorylation status of RPRM was crucial to the nuclear translocation of RPRM and its regulation on ATM, we also explored how RPRM phosphorylation affected cellular sensitivity to DNA damage. As shown in Figure 6i and j, overexpressing PRRM-S98A mutant alone caused a greater reduction in colony formation and a greater increase in background micronucleus frequency in H460 cells than overexpressing full-length RPRM did. Moreover, H460 cells with PRRM-S98A mutant expression showed much greater sensitivity to DNA damaging agents such as IR (Fig. 6i, j) and cisplatin (Fig. 6k). Furthermore, the radiosensitivity of H460 cells was enhanced when RPRM phosphorylation was suppressed by CDK4/6 inhibitor, Palbociclib, and this increase was more significant when RPRM was over-expressed (Fig. 6l, m). To further confirm the result that inhibition of RPRM phosphorylation confers cellular sensitivity to DNA damage, using RPRM^-/-^ MEFs re-expressing full-length RPRM, RPRM-S98A and ΔRPRM(79-109) mutants, we found that compared with the negative control RPRM^-/-^ MEFs and RPRM^-/-^ MEFs re-expressing full-length RPRM, the RPRM^-/-^ MEFs re-expressing PRRM-S98A mutant exhibited higher background and radiation-induced micronucleus frequency, suggesting that inhibition of RPRM phosphorylation enhanced the sensitivity of normal cells to spontaneous and IR-induced DNA damage (Fig. 6n). It was also worth noting that re-expressing ΔRPRM(79-109) mutants only caused a slight but significant increase in micronucleus formation post IR when compared with NC (Fig. 6n), suggesting that a fully intact C-terminal end of RPRM was also important to the radiosensitizing effect of RPRM in addition to its unphosphorylation status. These data indicated that for both cancer and normal cells, inhibition of RPRM phosphorylation did confer cellular sensitivity to DNA damage. This agreed with the negative regulatory effect of RPRM on ATM involving the RPRM phosphorylation mediated by CDK4/6 and ATM nuclear export. It also provided a novel mechanism for the radiosensitizing effect of Palbociclib for cancer cells.^38–40^ These results suggest that RPRM may serve as a potential target for cancer therapy and radiation protection.

### RPRM is a potential target for cancer therapy and radiation protection

To further confirm the determining role of RPRM in radiosensitivity, we performed xenograft assay, and found that RPRM overexpression not only significantly inhibited tumor growth, which agreed with its tumor suppressor characteristics,^21–24^ but also sensitized tumors to radiotherapy (Fig. 7a, b). With Hematoxylin & Eosin (H&E) staining, we observed that the tumors formed by H460-RPRM cells grew in loose clusters and showed a feature of high differentiation, pleomorphism, extensive keratinzation and spontaneous apoptosis. In contrast, the tumors formed by H460-NC cells exhibited a feature of strong basophilia, low differentiation and keratinization (Fig. 7c). Moreover, IR exposure enhanced apoptosis more substantially in H460-RPRM cell-formed tumors than in H460-NC cell-formed tumors (Fig. 7c).

**Figure 7.**
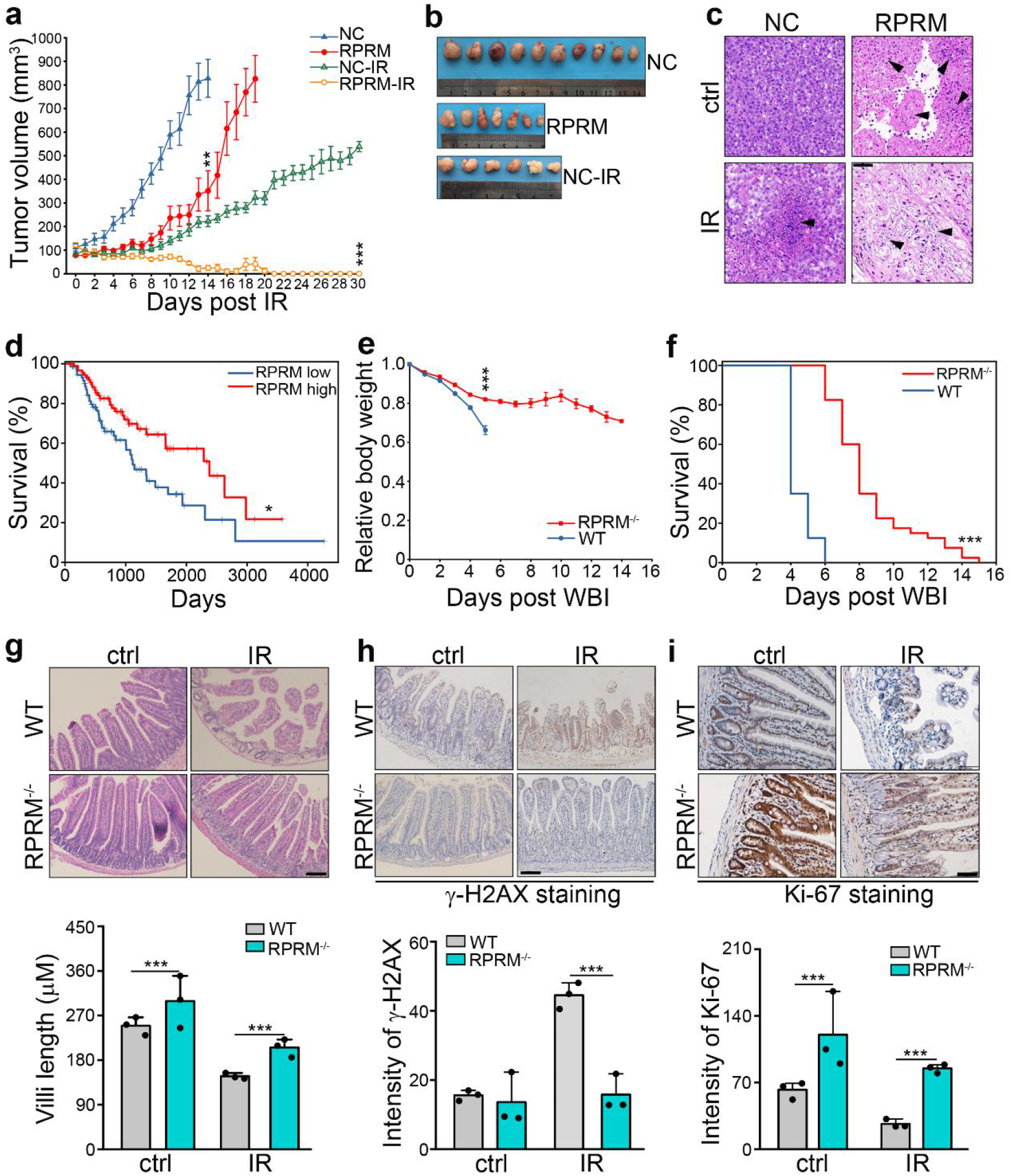
RPRM is a potential target for cancer therapy and radiation protection. **a, b** Comparison of the tumor growth of the four groups after a single X-irradiation dose of 5 Gy. Tumor volumes were calculated by the formula: volume = a (larger diameter) b^2^ (smaller diameter)/2 (mice: *n* = 10 for NC, *n* = 7 for RPRM, *n* = 6 for NC+IR, *n* = 5 for RPRM+IR). Data are presented as mean ± SEM, two-way ANOVA test followed by Tukey’s correction was performed (**a**) and representative pictures of tumors (**b**) excised on day 14 (NC), 18 (RPRM), 30 (NC-IR), respectively. No pictures of irradiated tumors with RPRM overexpression were provided since the rumors shrank completely by the end of observation. **c** H&E staining of the tumor sections showing that RPRM overexpression caused more severe cell death. Scale bar, 50 μm. Black arrows indicate apoptosis of tumor cells. **d** The cancer genome atlas (TCGA) lung squamous cell carcinoma data set shows that the clinical outcome after radiotherapy (10-y overall survival) was associated with the expression of RPRM. Kaplan-Meier survival curves and Log rank statistics are shown. **e** Normalized body weight changes of RPRM^-/-^ and WT mice after receiving 9 Gy WBI showed a more dramatic decline in the body weights of WT mice than in those of RPRM^-/-^ mice. **f** Kaplan-Meier analysis of the survival of WT and RPRM knockout mice after receiving 9 Gy WBI (*n* = 40 for each group including 20 male and 20 female mice). **g** H&E staining of the small intestines of RPRM^-/-^ and WT mice on day 3.5 after 9 Gy WAI. Villi length was quantified using CellSens Standard 1.8.1 (*n* = 3 for each group). Scale bar, 100 μm. **h** Immunohistochemistry staining of γ-H2AX on day 3.5 after 9 Gy WAI. Positive cells were counted by using image pro plus 6.0 (*n* = 3 for each group). Scale bar, 100 μm. **i** Immunohistochemistry staining of Ki-67 on day 3.5 after 9 Gy WAI. Positive cells were counted by using image pro plus 6.0 (*n* = 3 for each group). Scale bar, 50 μm. \**P* < 0.05; \*\**P* < 0.01; \*\*\**P* < 0.001.

It was reported that RPRM is expressed at a low level in a variety of cancers due to hypermethylation of RPRM promoter. ^22–25^ To further determine the therapeutic relevance of RPRM in cancers, we performed TCGA analysis. According to the TCGA lung squamous cell cancer database, the good prognosis of lung cancer patients after radiotherapy was associated with high RPRM mRNA expression (Fig. 7d). The same tendency was observed in glioma patients (Supplementary Fig. S10a). Cervical and endocervical cancer patients showed similar trend, although the difference between patients with high and low RPRM expression was not statistically significant (Supplementary Fig. S10b). All these data indicate that RPRM is a potential therapeutic target for cancers.

Furthermore, we investigated the radiosensitivity of RPRM knockout mice. RPRM knockout mice appeared normal and were indistinguishable from their WT littermates (Supplementary Fig. S11a, b). After whole body irradiation (WBI), compared with WT mice that lost weight sharply and all died within 6 days, RPRM knockout mice lost less weight and even made a slight improvement 7 days after WBI before the second wave of weight loss starting on day 11, and eventually all died within 15 days post WBI (Fig. 7e, f). Both male and female RPRM knockout mice showed similar results, although it seemed that RPRM knockout had better protective effect on male mice than on female mice (Supplementary Fig. S11c, d). These data indicated that RPRM knockout significantly delayed mouse death after exposed to WBI. Moreover, we examined radiation-induced intestinal injury in RPRM knockout mice. After whole abdominal irradiation (WAI), compared with WT mice, RPRM knockout mice showed much less severe villus injury (Fig. 7g), less DNA damage (γ-H2AX) (Fig. 7h) and more cell proliferation (Ki67) (Fig. 7i), indicating that RPRM knockout dramatically protected mice against radiation-induced intestinal injury. All these results indicated that RPRM deficiency protected mice from IR-induced DNA damage, thus may serve as a target for radiation protection. Taken together, it was confirmed in vivo that RPRM is a potential target for cancer treatment and radiation protection.

## Discussion

Here, we discovered that RPRM negatively regulates ATM levels and inhibits DNA damage repair, thus enhancing cellular sensitivity to genotoxic agents. After induced by DSBs, RPRM rapidly translocates from cytoplasm to nucleus, in which CDK4/6 is involved through mediating RPRM phosphorylation at serine 98, interacts with ATM and promotes the nuclear export and proteasomal degradation of ATM resulting in a reduction of ATM levels, thus inhibits DNA repair pathway. Therefore, RPRM, this poorly understood protein, is actually an important novel regulator of cellular sensitivity to DNA damage.

Despite the intensive investigation into the vital role of ATM as an apical conductor of DDR relaying a wide-spread signal to a variety of downstream effectors in response to DNA DSBs,^2–5, 9^ the molecular mechanisms underlying how ATM itself is regulated, in particular shortly after DNA damage induction, remain to be elucidated. In this study, we revealed that RPRM is a novel negative regulator of ATM. We found that ATM expression levels declined conspicuously in RPRM-overexpressed cells soon after the induction of DNA DSBs. Moreover, RPRM modulated the down-regulation of ATM at protein level but not at mRNA level. More specifically, RPRM down-regulated ATM protein by promoting its nuclear export and proteasomal degradation. Interestingly, compared with NC cells, the cells overexpressing RPRM showed a higher basal level of cytoplasmic ATM, and the level substantially increased post IR, which was almost reversed by using leptomycin B, a potent inhibitor of the nuclear export of proteins. It has been previously shown that exposure to IR or etoposide induces the nuclear-cytoplasmic translocation of activated ATM.^41^ However, we did not observe any increase in the phosphorylated ATM levels in the cytosol of the cells overexpressing RPRM. These results indicated that RPRM promoted the nuclear-cytoplasmic translocation of ATM but not that of activated ATM. In addition, it suggested that the nuclear-cytoplasmic translocation of ATM was different from that of activated ATM in terms of the consequences. While the nuclear export of activated ATM triggered NF-κB activation,^41^ the nuclear export of ATM appeared to lead to its degradation and down-regulation, suggesting that the degradation and down-regulation of ATM were associated with its cytoplasmic localization. This agrees with what was previously reported that the total ATM protein level is positively correlated with its nuclear localization and ATM underexpression is linked to its abnormal cytoplasmic localization.^30^ Our results also provide a potential mechanism for ATM down-regulation, i.e. degradation due to its abnormal cytoplasmic localization mediated by RPRM. Interestingly, in addition to the canonical function of ATM in DNA damage repair signaling pathway, which is dependent on its nuclear localization, accumulating evidence shows that cytoplasmic ATM may play some roles in Akt activation, autophagy in response to oxidative stress, etc.^42, 43^ This study suggests that the ATM in RPRM-overexpressed cells containing higher cytoplasmic ATM levels may function differently compared with that in the parental cells. Giving our finding that RPRM negatively regulates ATM through changing its subcellular localization and protein levels, further studies are warranted on how RPRM modulates ATM translocation and degradation, as well as what functions the cytoplasmic ATM may have when RPRM is overexpressed.

RPRM has been proposed as a cytoplasmic protein since it was identified.^20^ However, in the present study, we discovered that RPRM translocated to nucleus shortly after IR treatment and bound to ATM, indicating that in spite of a cytoplasmic protein, RPRM can translocate rapidly from cytosol to nucleus under DNA damage stress. The function of a protein is usually associated with its subcellular localization. Our finding on the change in the subcellular localization of RPRM upon DNA damage indicates that RPRM has some novel functions related to its nuclear translocation, e.g. its negative regulatory effects on ATM and DNA repair. We further found that the nuclear transport of RPRM was governed by CDK4/6-mediated phosphorylation of RPRM at serine 98. Inhibition of CDK4/6 by Palbociclib promoted the nuclear translocation of RPRM, mutation or deletion of serine 98 of RPRM protein also facilitated its nuclear transport, which was very likely due to its unphosphorylation and stabilization. This is similar to the reports that site-specific phosphorylation of proteins such as p53 and neurofibromatosis-2 (NF2) leads to their degradation by ubiquitination.^44, 45^ As a result of the unphosphorylation of RPRM and its nuclear translocation mediated by CDK4/6 inhibition, ATM was exported from nucleus and down-regulated. There is some evidence suggesting that ATM may be associated with CDK4. It was found that the E3 ligase WSB1 promoted ATM ubiquitination and degradation after its activation through CDK4-mediated phosphorylation.^46^ Our results establish a novel connection between CDK4/6 and ATM, which is that CDK4/6 indirectly modulates ATM levels through RPRM. We also provide another potential mechanism for the inhibition of ATM activation by CDK4/6 inhibitor Palbociclib.^38^

The previous studies have suggested that RPRM may be involved in cell cycle arrest and apoptosis induced by DNA damage.^20, 23^ But whether it participates DNA repair pathway remain unknown. In this study, in agreement with its nuclear translocation upon DNA damage as we described above, we clearly demonstrated that RPRM played an important novel role in DNA DSB repair. We found that RPRM overexpression inhibited both HR and NHEJ pathways resulting in impaired DNA damage repair and more DSB accumulation, in turn increased cellular radiosensitivity. This may be associated with its negative regulatory effect on ATM. These results were consistent with the hypersensitivity of ATM-mutated cancer cells and A-T cells to IR.^47, 48^ Inhibiting DDR pathway is one of the important mechanisms underlying the capability of sensitizers to sensitize cancer cells to genotoxic agents.^49^ Thus, proteins that play critical roles in DDR such as ATM, CDK4/6, etc. have been proposed as important therapeutic targets for cancer therapy. Several ATM inhibitors and CDK4/6 inhibitors have been demonstrated to exhibit a synergistic effect with radiotherapy both in vitro and in vivo.^15–17, 38–40^ Early clinical development has also been initiated to explore the implication of some ATM inhibitors.^19^ Here we identified RPRM as a critical protein in DNA damage repair pathways. We also confirmed its inhibitory effect on tumor growth, which is in agreement with its tumor suppressor properties.^21, 22^ These results indicate that RPRM may serve as a novel target for cancer therapy. Furthermore, it is worth pointing out that RPRM may inhibit DNA repair through other targets in addition to ATM. Since the phosphorylation of 53BP1 and the focus formation may involve ATM, ATR and DNA-PKcs,^50^ our results showing an inhibited 53BP1 foci formation when RPRM was overexpressed did not rule out the possibility of the regulatory effect of RPRM on ATR and DNA-PKcs. As a matter of fact, our unpublished results showed that DNA-PKcs was also down-regulated in cells with RPRM overexpression (Unpublished data).

In addition to enhancing cancer cell killing in cancer therapy, normal tissue must be protected against therapy-induced damage. However, until now radiation injury caused by radiotherapy and radiation accident remains a huge challenge. Short-lived free radicals produced by IR play a big role in radiation injury due to their damaging effects on cellular components especially DNA.^51^ Therefore, free radicals are a common target for radiation protection. In fact, a majority of radioprotectors tested so far are antioxidants that scavenge free radicals thus alleviating cellular damage caused by IR.^51^ However, despite extensive testing, to date there is only one radioprotector, i.e. amifostine, that was approved by FDA for clinical use, and its use is very limited due to its inconclusive protective effects in clinical trials, its side effects (nausea and hypotension) and delivery modality (it must be delivered prior to irradiation).^52^ This suggests that targets other than free radicals need to be explored in order to develop better and more effective radioprotectors. In this study, in addition to cancer cells, we further confirmed the determining role of RPRM in the radiosensitivity of normal cells and tissues. We found that RPRM-deficient cells were more resistant to IR than their WT counterparts. More intriguingly, RPRM-knockout mice exhibited much more delayed death after exposed to WBI and much less severe intestinal injury after exposed to WAI compared with WT mice. Giving that RPRM overexpression increases IR-induced DNA damage and impairs DNA repair, these results suggest that RPRM, a novel important DDR protein, may be also a potential target for radiation protection.

In summary, we discovered a novel critical role of RPRM in DNA repair and cellular sensitivity to DNA damage, and revealed an important underlying mechanism that RPRM translocates to the nucleus and interacts with ATM to promote the nuclear-cytoplasmic translocation of ATM resulting in its degradation and down-regulation, in which the phosphorylation of RPRM at serine 98 mediated by CDK4/6 is involved. These data not only shed light on the vital regulatory effect of RPRM on ATM, but also highlight the potential implications of RPRM in both cancer therapy and radiation protection. Additionally, active investigation is ongoing to characterize this poorly understood protein, RPRM, and explore its other functions.

## Materials and Methods

### Mouse experiments

All animal procedures were approved by the Ethics Committee of Soochow University. Soochow University Medical Experimental Animal Care Guidelines in accordance with the National Animal Ethical Policies of China were strictly adhered to in all animal experiments. We established RPRM^-/-^ C57BL/6 mice, which were processed by GemPharmatech Co., Ltd under commission. The WT C57BL/6 mice obtained during breeding were used as control. Male BALB/c nude mice (3-5 weeks) were purchased from the Experimental Animal Centre of Soochow University. All mice were specific pathogen free (SPF), and were kept in SPF animal facility of Soochow University Experimental Animal Centre. All mice were randomly divided into different experimental groups. For the survival experiment, RPRM^-/-^ C57BL/6 mice and WT mice aged 3 months (*n* = 40/group, including both male and female) were irradiated with 9 Gy whole body X-irradiation (2.0 Gy/min, 320 kVp, X-RAD320ix, PXi). The body weights of mice were measured daily until their deaths. For radiation-induced intestinal injury, male RPRM^-/-^ C57BL/6 mice and WT mice aged 3 months (*n* = 3/group) received 9 Gy of whole abdominal X-irradiation (WAI, 1.16 Gy/min, 160 kVp, RAD SOURCERS2000 X-ray machine). Mice were sacrificed on day 3.5 after WAI, and the small intestines were harvested and processed for immunohistochemistry and HE staining. For xenograft cancer experiment, H460-NC (1×10^6^) and H460-RPRM (3×10^6^) cells were subcutaneously injected into the right flank of BALB/c nude mice. Once reaching approximate 100 mm^3^ in volume, the tumors were topically irradiated with 5 Gy X-rays (1.16 Gy/min, 160 kVp, RAD SOURCERS2000 X-ray machine). And the tumor growth was then monitored daily using caliper measuring. Tumor volumes were calculated as length×width^2^/2. Mice were sacrificed when tumor volume reached 1000±100 mm^3^ or on day 30 post irradiation.

### Cell lines, cell culture and reagents

H460 and A549 human non-small cell lung cancer cell lines, SNU-1 human gastric carcinoma cell line and AGS human gastric adenocarcinoma cell line were purchased from the Cell Bank of Chinese Academy of Sciences. H460 and SNU-1 cells were cultured in RPMI1640 (Sigma-Aldrich) containing 10% FBS (Hyclone), 2.5 g/l glucose (Sinopharm Chemical Reagent Co., Ltd), 1.5 g/l sodium bicarbonate (Sinopharm Chemical Reagent Co., Ltd), 1% sodium pyruvate solution (Sigma-Aldrich) and 1% penicillin-streptomycine (Beyotime). A549 and AGS cells were cultured in F12K (Sigma-Aldrich) with 10% FBS, 1% penicillin-streptomycine and 2.5 g/l sodium bicarbonate. WS1 human embryonic dermal fibroblasts were purchased from American Type Culture Collection (ATCC). HaCaT human immortalized keratinocytes were obtained from China Center for Type Culture Collection. Both cell lines were cultured in DMEM medium with high glucose (Gibco) containing 10% FBS and 1% penicillin-streptomycine. All cell lines were tested negative for mycoplasma. MEFs were isolated as described previously.^46^ Briefly, mouse embryos were harvested at E13 and the internal organs were removed, then the embryo bodies were digested using 0.25% trypsin-EDTA (Beyotime) for 30 min at 37 °C, and neutralized with DMEM containing 10% FBS. The tissue homogenate was then filtered through 40 μm nylon cell strainer (Falcon) and centrifuged at 1000 rpm for 5 min, and the pellets were resuspended and cultured in completed DMEM.

Inhibitors: Cycloheximide (CHX, Selleck; 0.1 mg/ml) for protein synthesis inhibition; Leptomycin B (LMB, Beyotime; 10 nM) for nuclear export inhibition; Palbociclib (Palb, Selleck; 50 μM) for CDK4/6 inhibition and MG132 (Selleck; 10 μM) for proteasome inhibition; Cisplatin (CDDP, Sigma-Aldrich; 15 μM) for inducing DNA damage in tumor cells.

### Plasmids and stable transfection

pCMV3-N-HA negative control vector and human RPRM plasmids (full length, S98A mutant and Δ(79-109) mutant) were purchased from Sino Biological Inc. The RPRM and negative control plasmids were transfected into cells using Lipofectamine™ 2000 (Invitrogen). The medium was replaced with fresh complete medium 6 h later. Cells were subcultured 48-72 h after transfection and were grown in selective medium containing 8 μg/ml puromycin (Beyotime). Single-cell clones were selected and multiplied in growth medium containing 4 μg/ml puromycin.

### Lentiviral infection

Lentiviral constructs (pHBLV for shRNA) were purchased from Hanbio Biotechnology Co., LTD. 7×10^5^ cells were cultured in 1 ml medium containing viral supernatant in 6-well plates for 4 h at 37 °C, and were added with 1 ml fresh medium and continued to culture for 24 h. Gene expression and cell viability were assessed at least 48 h post-infection. shRNA sequences are listed in Supplementary Table S1.

### Neutral comet assay

Cells were collected after irradiation and resuspended in cold PBS at a density of 1×10^5^ cells/ml. 50 μl of cell suspension was combined with 500 μl of molten LMAgarose (Trevigen), and 50 μl of mixture was pipetted onto CometSlide™ (Trevigen^®^) immediately according to the manufacture’s instruction. In brief, slides were placed at 4 °C in the dark for 10 min and then sequentially immersed in lysis solution and 1×neutral electrophoresis buffer for 1 h and 30 min, respectively, followed by electrophoresis at 17 V for 45 min at 4 °C (Mini-Sub^®^ Cell GT, BIORAD). Slides were transferred to DNA precipitation solution for 30 min at room temperature (RT). After immersion in 70% ethanol for 30 min, slides were dried at 37 °C for 15 min. Then slides were stained in 1×SYBR^®^ Gold (Thermo Fisher Scientific) staining solution for 30 min at RT in the dark. Slides were examined under a laser scanning confocal microscopes (FV2000, Olympus) with an excitation wavelength of 496 nm and an emission wavelength of 522 nm.

### RNA extraction and real-time RT-PCR

Total RNA was isolated using TRIzol^®^ Reagent (Life Technologies™) and quantified by NanoDrop 2000c (Thermo). According to the manufacture’s protocols, reverse transcription and real-time qPCR were performed using 5×All-In-One RT MasterMix (Abm) and EvaGreen 2×qPCR MasterMix (Abm), respectively. The primer sequences (Invitrogen) used for quantitative RT-PCR are listed in Supplementary Table S2.

### Protein extraction, co-immunoprecipitation and western blotting

For total protein extraction, cells were lysed in RIPA (Solibol) containing protease inhibitors (cOmplete ULTRA Tablets, Roche) for 1 h on ice, then centrifuged at 16,000 g for 10 min at 4 °C, and the supernatant was collected. For co-immunoprecipitation (IP), the supernatant was successively incubated with primary antibody of HA (AH158, Beyotime; 1:50) or ATM (GTX70103, GeneTex; 1:80) at 4 °C overnight and protein G-Agarose (Roche) at 4 °C for 3 h. After washing 5 times with RIPA, co-purified proteins by IP were analyzed by western blotting. Nuclear and cytoplasmic protein extraction was performed as previously described.^53^ For western blotting, after denature, protein extracts were separated on a 6%-12% SDS-PAGE gel and transferred to PVDF membrane. Following blocking with 5% bovine serum albumin (BSA, Beyotime) or 5% non-fat milk (Sangon Biotech), the proteins of interest were probed by sequential incubation with primary antibodies: HA (AH158, Beyotime; 1:1000), RPRM (GTX110976, GeneTex; 1:1000), Phospho-ATM (13050, CST; 1:1000), ATM (GTX70103, GeneTex; 1:1000), CDK4 (ab199728, Abcam, 1:1000), CDK6 (AC256, Beyotime; 1:1000) Histone H3 (GTX122148, GeneTex; 1:1000), LMNB2 (AF0219, Beyotime; 1:1000), β-actin (AA128, Beyotime; 1:1000), GAPDH (AF1186, Beyotime; 1:1000), and Tubulin (AT819, Beyotime; 1:1000) at 4 °C overnight and secondary antibodies (HRP-conjugated anti-rabbit/mouse IgG, Beyotime; 1:1000) at RT for 2 h, and detected on a FluorChem™ M System (Alpha) after treatment with ECL-plus (Beyotime).

### Immunofluorescence staining

Cells on coverslips were fixed in 4% paraformaldehyde (Sangon Biotech) for 15 min at RT. After permeabilization and blocking, cells were sequentially incubated with primary antibodies: γ-H2AX (ab22551, Abcam; 1:250), 53BP1 (ab21083, Abcam; 1:200), RAD51 (GTX100469, GeneTex; 1:1000), ATM (GTX70103, GeneTex; 1:200), phospho-ATM (SAB1413106, Sigma-Aldrich; 1:100), HA (AF0039, Beyotime; 1:100) at RT for 45 min (γ-H2AX, 53BP1, RAD51) or 4 °C overnight (ATM, phospho-ATM and HA), and secondary antibodies: DyLight 488 (mouse IgG antibody, GTX213111-04, GeneTex; 1:500), Dylight 488 (rabbit IgG antibody, GTX213110-04, GeneTex; 1:500), Cy3 (mouse IgG antibody, S0012, Affinity; 1:500) and Cys (rabbit IgG antibody, S0011, Affinity; 1:500) at RT for 50-120 min, followed by counterstaining with 5 μg/ml DAPI (Beyotime) at RT for 5 min. Cells were then imaged under a Olympus FV1200 confocal fluorescence microscope.

### Colony formation assay

Cells were seeded in 60 mm dishes or 6-well plates at appropriate densities according to different radiation doses and cell lines. After irradiation cells were continuously cultured for 2 weeks, followed by fixation in methanol for 15 min at RT and staining with methylene Blue (Sangon Biotech). The colonies with more than 50 cells were counted, and the percentage of cell survival was calculated. Sensitization enhancement ratio (SER) was calculated as the ratio of the irradiation dose required for the control group to obtain 30% survival to the radiation dose required to obtain the same survival rate after overexpression of RPRM.

### Micronucleus formation assay

Cells seeded on coverslips were irradiated, 24 h later, cells were fixed with fixative (methanol: acetic acid/3:1 in volume) for 15 min at RT, followed by staining with DAPI. Cells were examined under a Leica DM2000 microscope using a 40× objective lens. Nuclei with micronucleus were counted as positive, 1000 nuclei were examined for each sample.

### Mass spectrum

Purified RPRM peptides (77-109, ChinaPeptides; 1 μg) were incubated with purified recombinant CDK4 (Origene; 0.2 μg) and CDK6 proteins (Origene; 0.2 μg), respectively. The reaction products were resolved in SDS-PAGE gel and the gel slices including all proteins were excised. After gel digestion, LC-MS/MS analysis was performed at the JingJie PTM Biolab Co. Ltd. Proteome Discoverer 1.3 was used to process the MS/MS data. Trypsin/P was used as cleavage enzyme. For precursor and fragment ions, mass error was set to 10 ppm and 0.02 Da, respectively. High peptide confidence was selected and peptide ion score was set > 20.

### H&E and immunohistochemistry staining

Tumors and mouse intestines were fixed in 4% paraformaldehyde for at least 24 h and dehydrated by gradient ethanol solution. Paraffin-embedded tissue sections were deparaffinized and rehydrated followed by staining with hematoxylin for 10 min and eosin for 1 min. For immunohistochemistry, antigen retrieval was performed in sodium citrate solution (pH 6.0) for 20 min at 98 °C after deparaffinization and rehydration. Then the sections were blocked in 10% goat serum containing 1% BSA for 2 h at RT and sequentially incubated with primary antibodies: γ-H2AX (ab22551, Abcam; 1:100) or Ki67 (GTX16667, GeneTex; 1:200) at 4 °C overnight and sencondary antibody (Zsbio) for 1 h at RT. After counterstained with hematoxylin, the sections were examined under a microscope (IX73, Olympus), and the images were analyzed using image pro plus 6.0.

### Statistics

All statistical analysis was performed using Origin 2019 software. Data are presented as the average of 3 independent experiments ± standard error (SEM). Statistical significance was calculated using two-sample t-test or two-way ANOVA with Tukey’s correction. Survival of mice and cancer patients was presented using Kaplan-Meier curves and significance was assessed using log-rank test. *P* < 0.05 was considered significantly different between groups.

## Acknowledgments

This work was supported by the National Natural Science Foundation of China (Grants U1632270 and U1967220) and the Priority Academic Program Development of Jiangsu Higher Education Institution (PARD).

## Author contributions

Conceptualization: H. Y.; Validation: J. W., J. C.; Formal analysis: Y. Z, G. O; Investigation: Y. Z, G. O, Z. Y, Z. Z, Q. C., M. L.; Writing: H. Y., Y. Z.; Visualization: Y. Z, G. O; Supervision: H. Y.; Project administration: H. Y., J. W., J. C.; Funding acquisition: H. Y, J. C.

