## Supplementary data for "RPRM negatively regulates ATM levels involving its phosphorylation mediated by CDK4/CDK6"

**This PDF file includes:**

Figures. S1 to S11
Tables S1 to S2

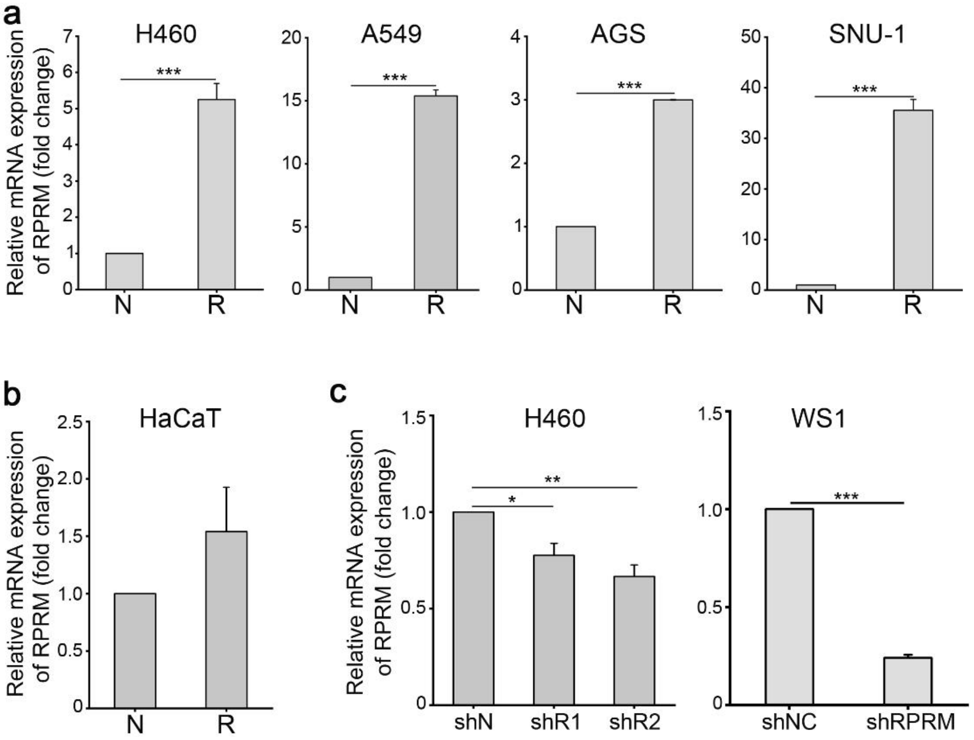


**Figure. S1.** Verification of RPRM-overexpression/knockdown in cancer and normal cells used in this study. **a, b** mRNA levels of RPRM in cancer cells (**a**) and HaCaT keratinocytes (**b**) transfected with RPRM (R) and NC (N) plasmids, respectively. **c** mRNA expression of RPRM in H460 and WS1 cells infected with lentiviral constructs containing shRNA targeting RPRM (shR/shRPRM) and NC shRNA (shN/shNC), respectively. Experiments were independently repeated three times with similar results. **P* <0.05; ***P* <0.01; ****P* <0.001.

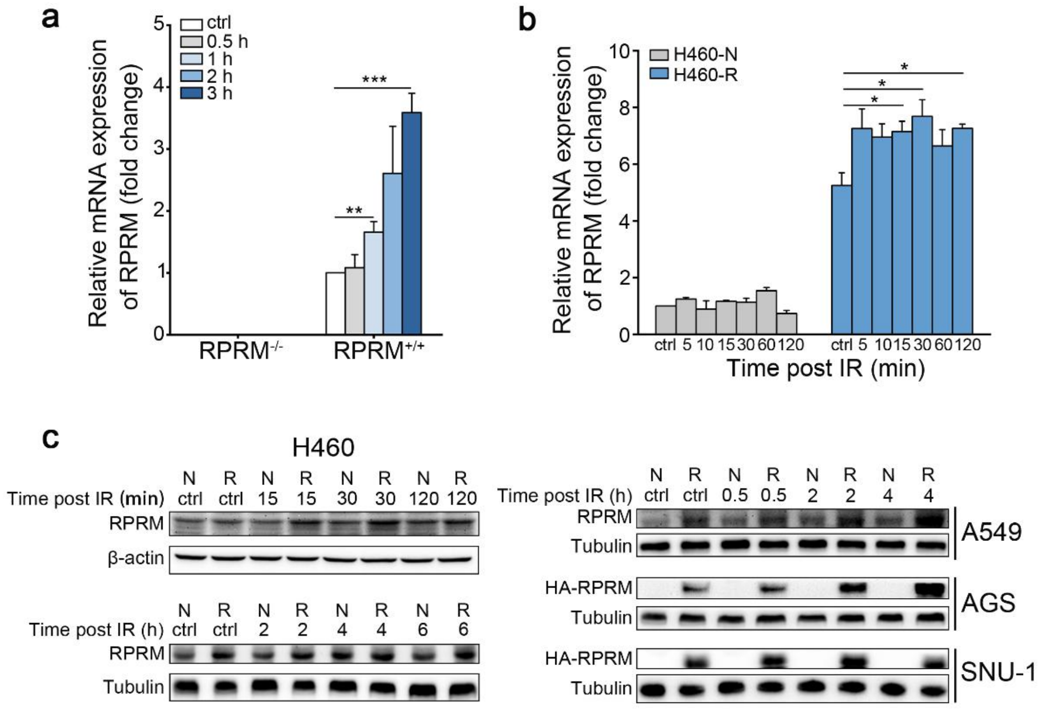


**Figure. S2.** Induction of RPRM in MEFs and cancer cells after X-irradiation. **a, b** Kinetics of the change in RPRM mRNA expression levels of $RPRM^{-/-}$ MEFs, $RPRM^{+/+}$ MEFs (**a**) and H460-NC (H460-N)/RPRM (H460-R) (**b**) after 20 and 2 Gy X-irradiation, respectively. **c** The change of RPRM protein levels in cancer cells overexpressed with RPRM after 2 Gy X-irradiation. Experiments were independently repeated three times with similar results. *$P$ < 0.05; **$P$ < 0.01; ***$P$ < 0.001.

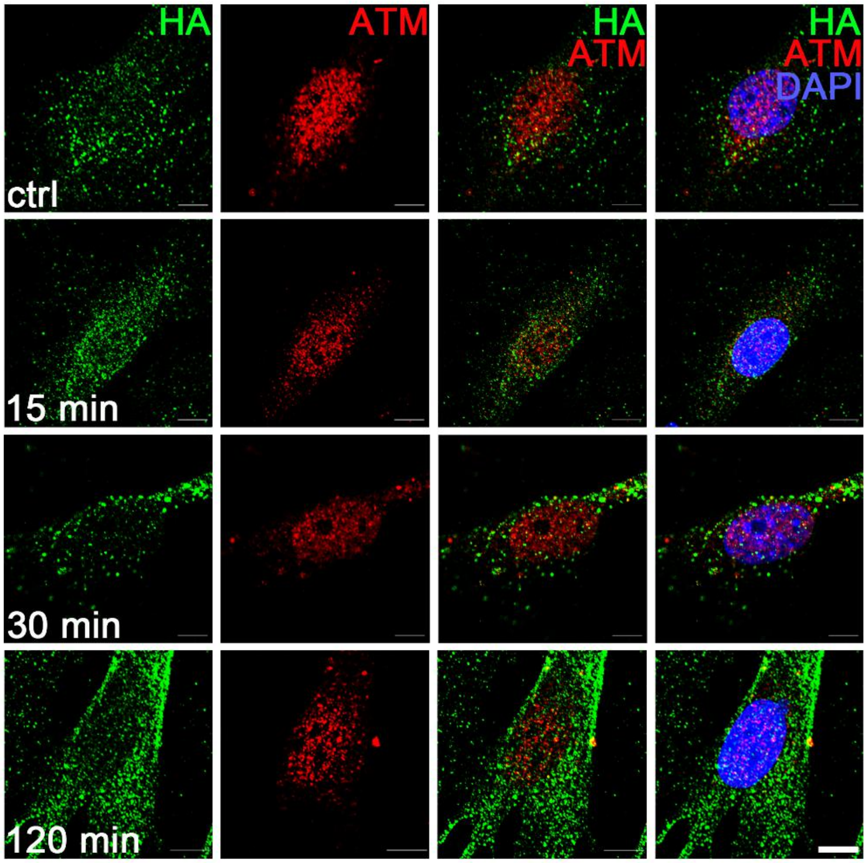


**Figure. S3.** RPRM binds to ATM. Representative immunofluorescence images showing that there was no co-localization of HA-tag and ATM in $RPRM^{-/-}$ MEFs transfected with negative control plasmids (NC) post 20 Gy X-irradiation. Scale bars, 10 μm. Experiments were independently repeated three times with similar results.

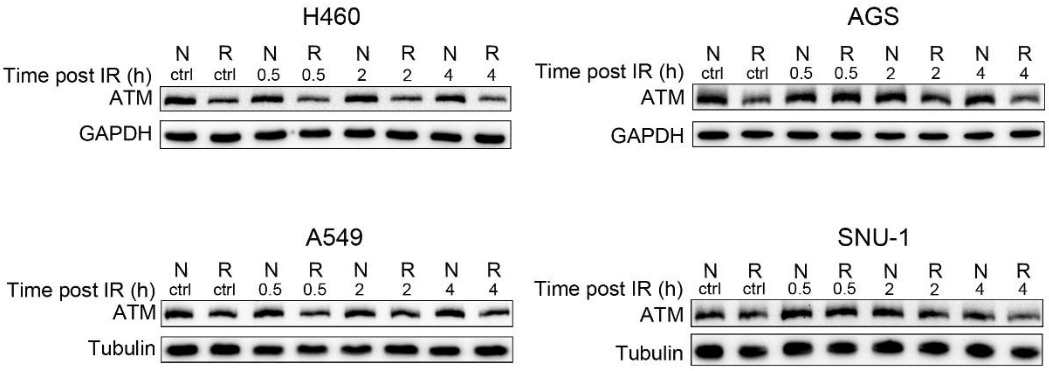


**Figure. S4.** RPRM down-regulates ATM levels. The ATM levels in cancer cells transfected with RPRM (R) plasmids were lower than those in negative control cells (N) post IR. Experiments were independently repeated three times with similar results.

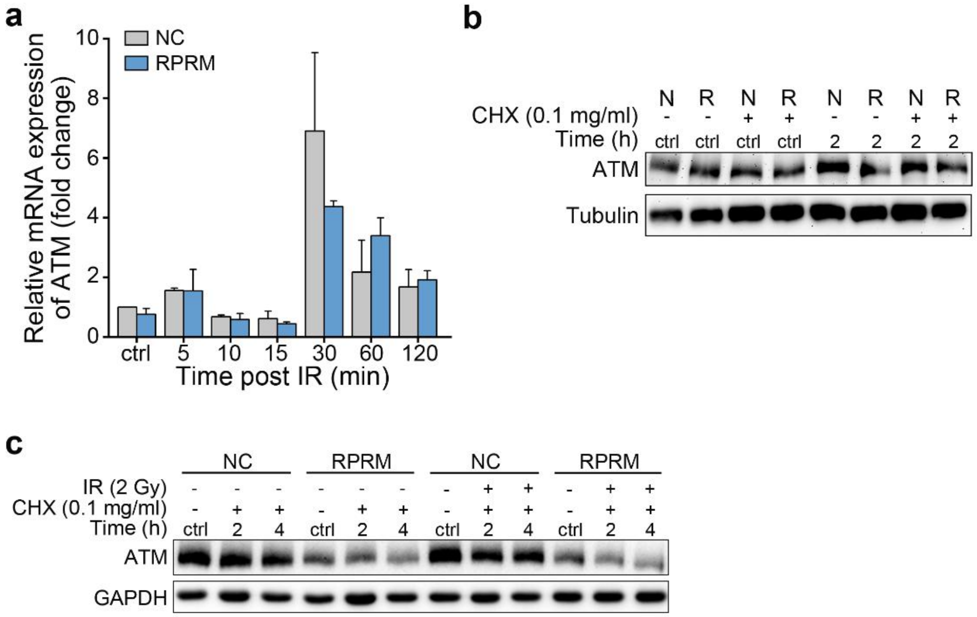


**Figure. S5.** RPRM promotes ATM degradation. **a** ATM mRNA expression levels of H460-NC/RPRM cells at different times after 2 Gy X-irradiation determined by qRT-PCR showed that RPRM overexpression did not significantly change the ATM mRNA levels upon IR when compared with the negative control cells. **b** Change in the ATM levels of H460-NC (N)/RPRM (R) cells 2 h after 2 Gy X-irradiation with and without cycloheximide (CHX, 0.1 mg/ml) pre-treatment. **c** Change in the ATM levels of AGS-NC/RPRM cells after exposed to 2 Gy X-rays with and without cycloheximide (CHX, 0.1 mg/ml) pretreatment. Experiments were independently repeated three times with similar results.

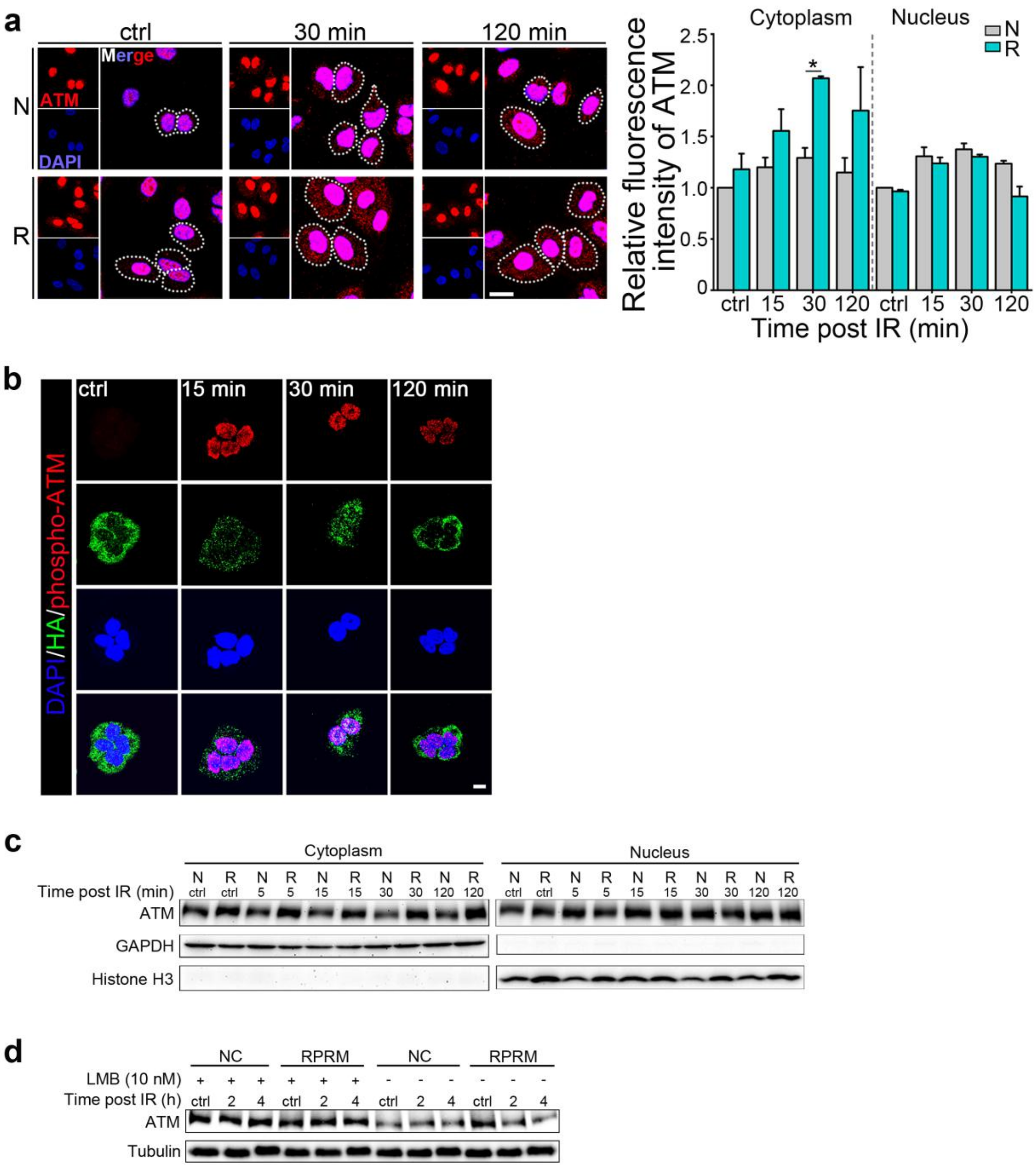


**Figure. S6.** RPRM affects the subcellular localization of ATM but not that of phospho-ATM, resulting in a reduction of total ATM levels. **a** Representative immunofluorescence images of ATM in H460-NC (N)/RPRM (R) cells at different times post 2 Gy X-irradiation and quantification of the fluorescence intensity of the nuclear and cytoplasmic ATM using image pro plus 6.0. Scale bar, 20 μm. White dotted circles indicate cytoplasmic ATM. **b** Phospho-ATM was still predominantly located in the nucleus of irradiated H460 cells overexpressing RPRM. Scale bar, 10 μm. **c** Western blotting images of the change in the cytoplasmic and nuclear ATM levels of H460-NC/RPRM cells at different times post 2 Gy X-irradiation. **d** Change of the total ATM levels in $RPRM^{-/-}$ MEFs transfected with NC/RPRM plasmids at different times post 20 Gy X-irradiation with or without leptomycin B (LMB, 10 nM) pretreatment. Experiments were independently repeated three times with similar results.

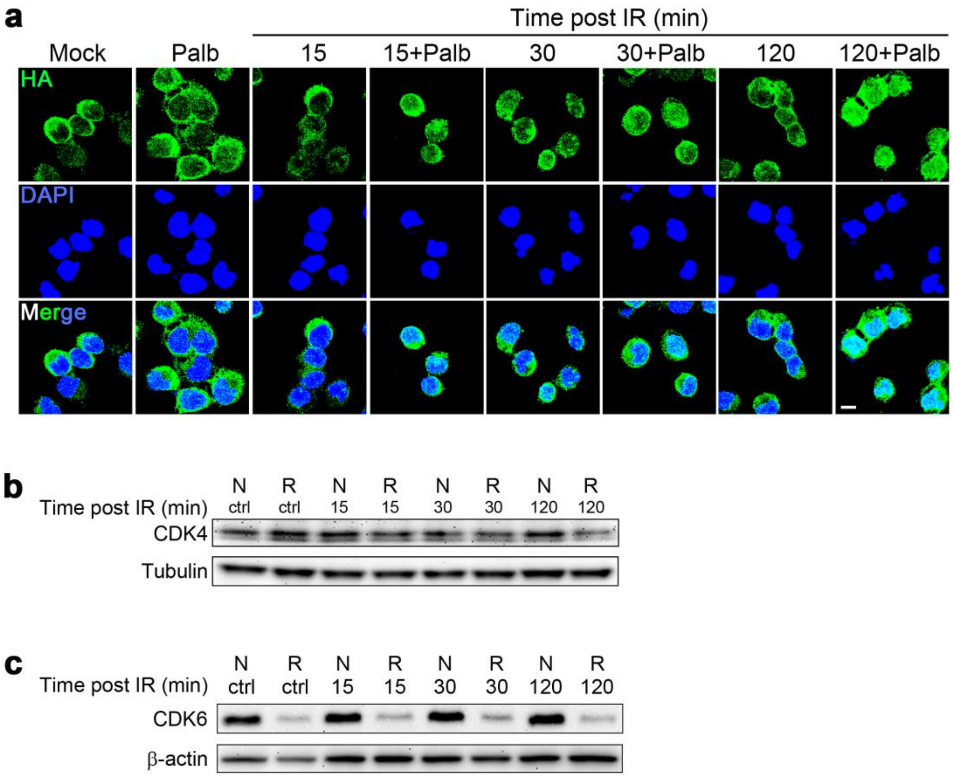


**Figure. S7.** The nuclear import of RPRM is enhanced by CDK4/6 inhibition. **a** Representative immunofluorescence images of H460-RPRM cells showing that the CDK4/6 inhibition by palbociclib pretreatment (Palb, 50 μM) for 1 h prior to 10 Gy X-irradiation enhanced the nuclear translocation of RPRM. Scale bar, 10 μm. **b, c** Change in the CDK4 (**b**) and CDK6 (**c**) levels of H460-NC (N)/RPRM (R) cells at different times after 2 Gy X-irradiation. Experiments were independently repeated three times with similar results.

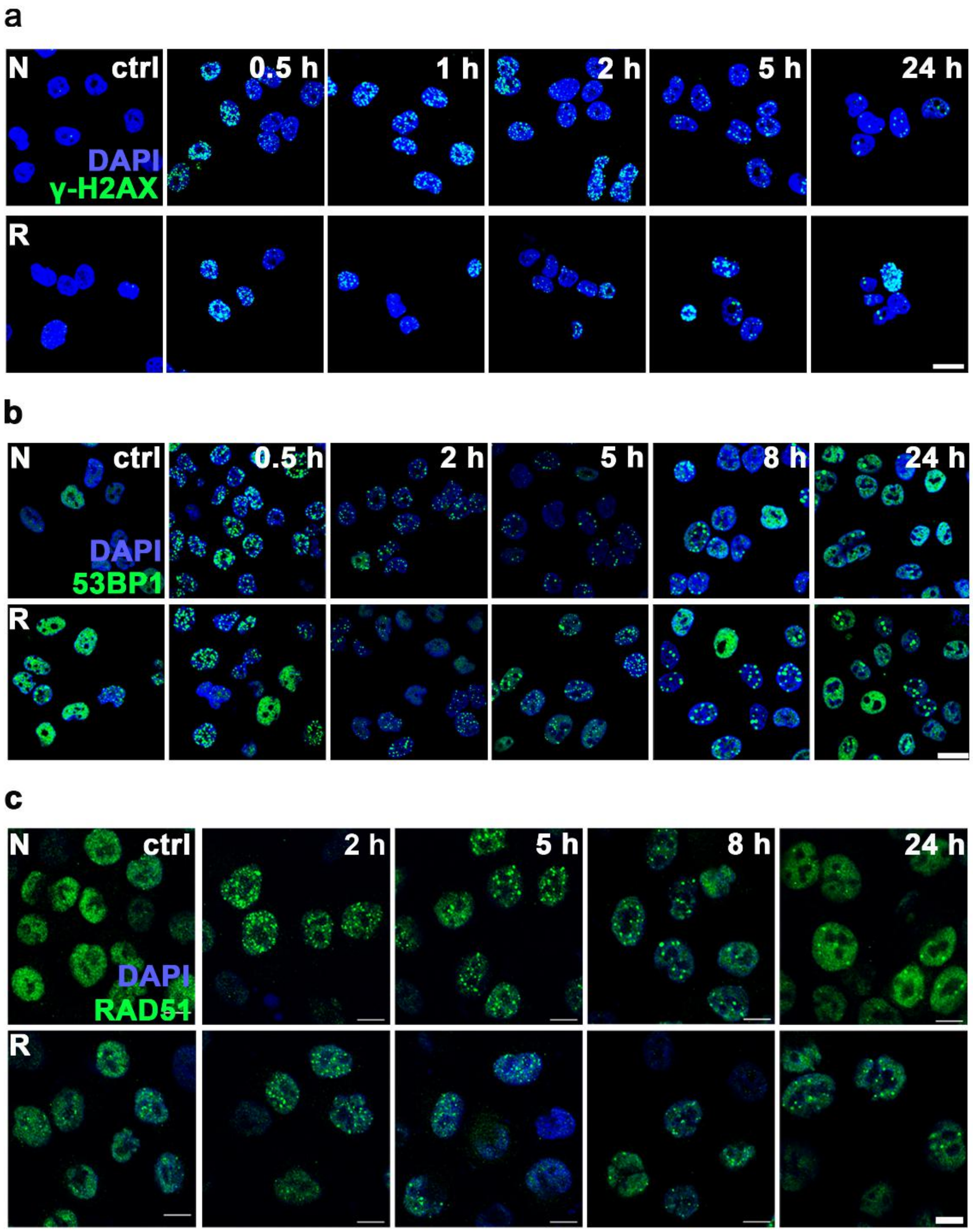


**Figure. S8.** RPRM inhibits DNA repair signaling pathway. Representative immunofluorescence images of γH2AX (**a** Scale bar, 20 μm), 53BP1 (**b** Scale bar, 20 μm) and RAD51 (**c** Scale bars, 10 μm) at the indicated time points post irradiation. H460-NC (N)/RPRM (R) cells were X-irradiated with 2 Gy and immunostained with antibodies to γH2AX, 53BP1 and RAD51, respectively. Experiments were independently repeated three times with similar results.

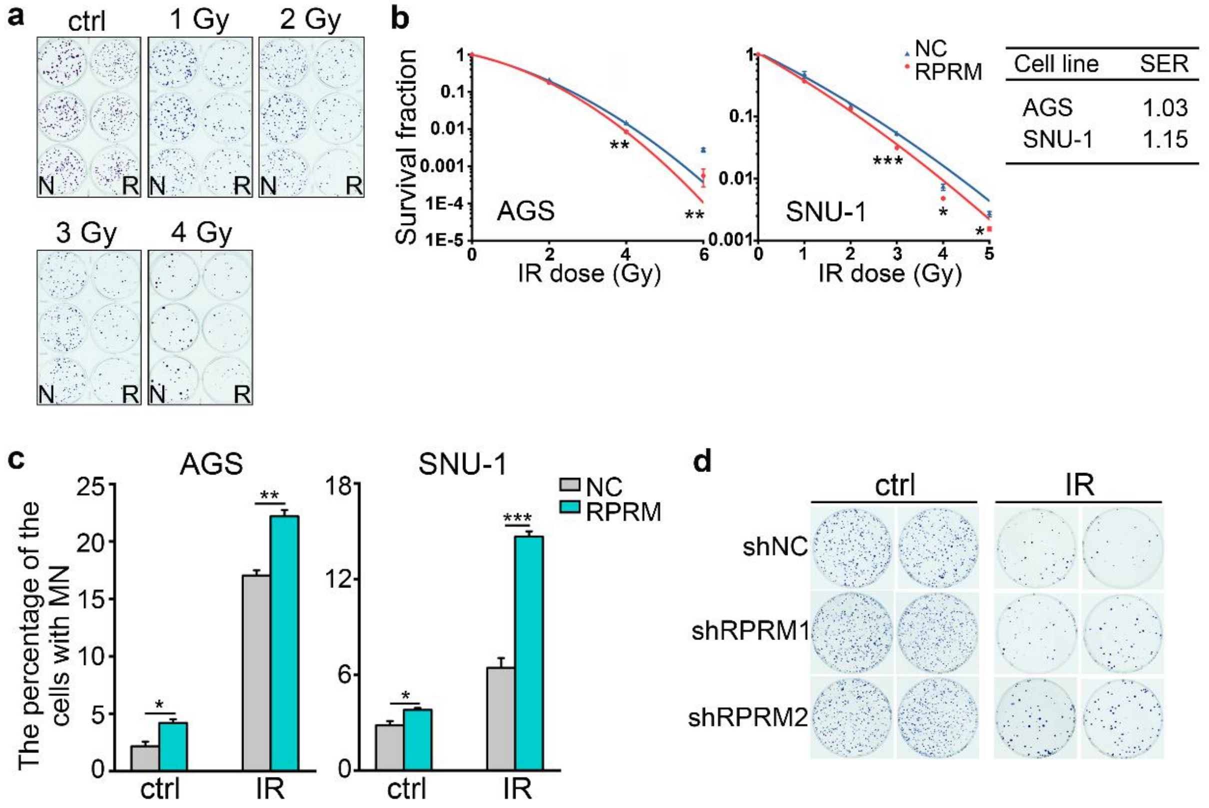


**Figure. S9.** RPRM enhances cellular radiosensitivity *in vitro*. **a** Representative pictures of H460 colonies after exposed to X-rays are showed. (N, negative control; R, RPRM overexpression) **b** Effect of RPRM overexpression on the survival of various cancer cell lines exposed to X-rays. Sensitization enhancement ratio (SER) was calculated according to the following formula: the radiation dose required for the control group to obtain 30% cell survival/the radiation dose required to obtain the same cell survival rate after overexpression of RPRM. **c** RPRM overexpression increased micronucleus formation in various cancer cells upon 2 Gy X-irradiation. **d** RPRM knockdown significantly increased cellular radioresistance manifesting as increased plating efficiency in irradiated H460 cells. Radiation dose was 4 Gy. Experiments were independently repeated three times with similar results. *$P$ < 0.05; **$P$ < 0.01; ***$P$ < 0.001.

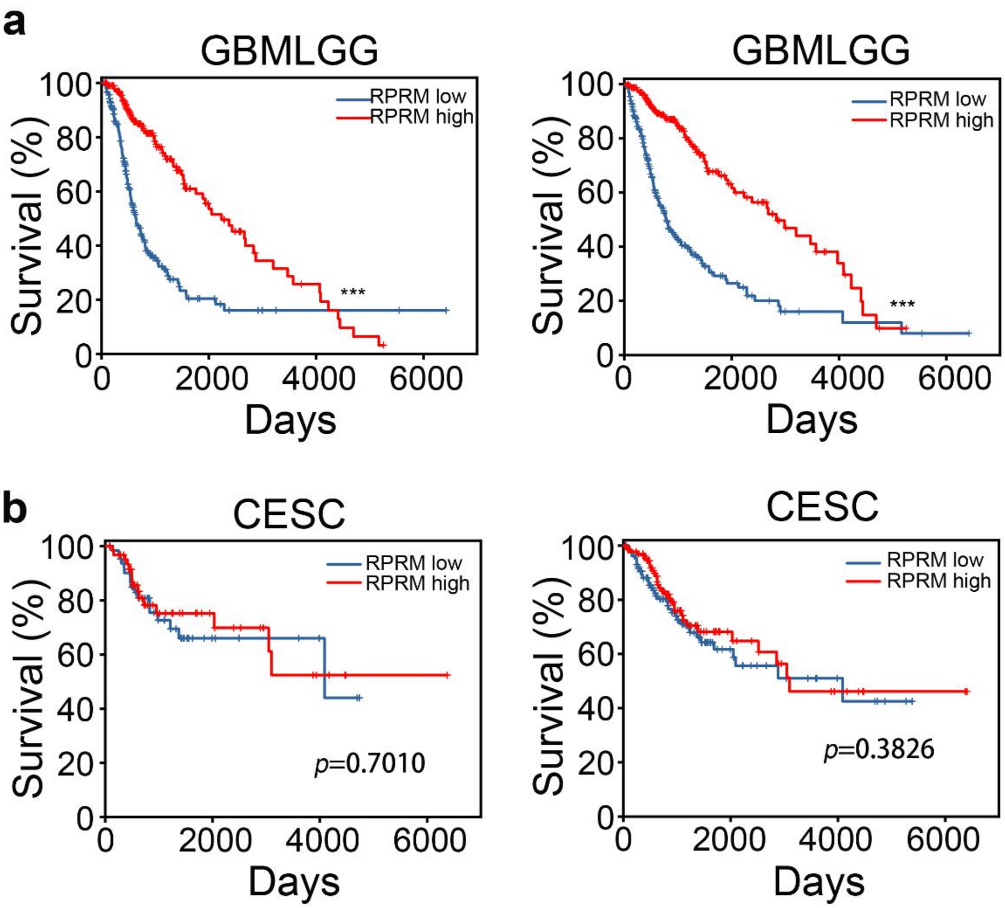


**Figure. S10.** The cancer genome atlas (TCGA) (**a** Glioma, GBMLGG; **b** cervical and endocervical cancer, CESC) data sets were used to determine that the clinical outcome with (left) or without (right) radiotherapy (10-y overall survival) was associated with the expression of RPRM. Kaplan-Meier survival curves and Log rank statistics are shown. ***$P < 0.001$.

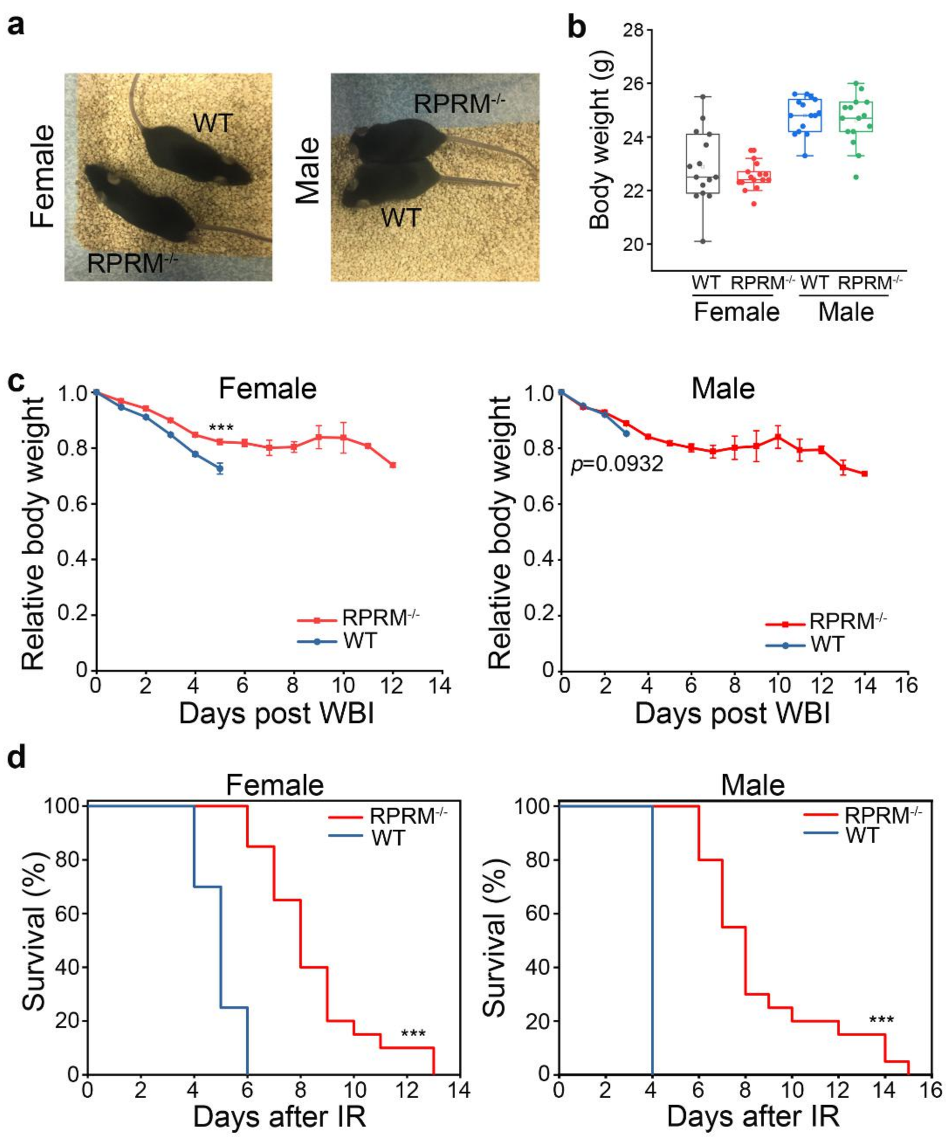


**Figure. S11.** RPRM knockout significantly reduced mouse radiosensitivity. **a** Representative pictures of 2-month-old RPRM$^{-/-}$ and WT mice. **b** Comparison of the body weights of 3-month-old RPRM$^{-/-}$ mice with those of WT mice showed that RPRM$^{-/-}$ mice were comparable to WT mice ($n$ = 15 for each group). **c** Normalized body weight changes of female and male RPRM$^{-/-}$ and WT mice after receiving 9 Gy WBI ($n$ = 20 for each group). **d** Kaplan-Meier analysis of the survival of female and male WT and RPRM knockout mice after receiving 9 Gy WBI ($n$ = 20 for each group). ***$P < 0.001$.

**Table S1. List of RPRM shRNA sequences.**

**Negative control top strand**

5'-GATCCGTTCTCCGAACGTGTCACGTAATTCAAGAGATTACGTGACACGTTCGGAGAATTTTTTC-3'

**Negative control bottom strand**

5'-AATTGAAAAAATTCTCCGAACGTGTCACGTAATCTCTTGAATTACGTGACACGTTCGGAGAACG-3'

**shRPRM1 top strand**

5'-GATCCGCATGATCAACTTCCTCGTGAATTCAAGAGATTCACGAGGAAGTTGATCATGTTTTTTG-3'

**shRPRM1 bottom strand**

5'-AATTCAAAAAACATGATCAACTTCCTCGTGAATCTCTTGAATTCACGAGGAAGTTGATCATGCG-3'

**shRPRM2 top strand**

5'-GATCCGTCATGTGCGTGCTCTCACTCATTCAAGAGATGAGTGAGAGCACGCACATGATTTTTTG-3'

**shRPRM2 bottom strand**

5'-AATTCAAAAAATCATGTGCGTGCTCTCACTCATCTCTTGAATGAGTGAGAGCACGCACATGACG-3'

**shRPRM3 top strand**

5'-GATCCGAGGGCATGATCAACTTCCTCGTGAATTCAAGAGATTCACGAGGAAGTTGATCATGCCCTTTTTTTG-3'

**shRPRM3 bottom strand**

5'-AATTCAAAAAAAGGGCATGATCAACTTCCTCGTGAATCTCTTGAATTCACGAGGAAGTTGATCATGCCCTCG-3'

**Table S2.** **List of primers used for qPCR.**

| Primer for qPCR | Sequence |
|---|---|
| Human RPRM forward | 5’-CTGGCCCTGGGACAAAGAC-3’ |
| Human RPRM reverse | 5’-TCAAAACGGTGTCACGGATGT-3’ |
| Human β-actin forward | 5’- AAAAGCCACCCCACTTCTCTCT-3’ |
| Human β-actin reverse | 5’- AATGCTATCACCTCCCCTGTGT-3’ |
| Human ATM forward | 5’-TGCAACAGGTCTTCCAGATG-3’ |
| Human ATM reverse | 5’-CAAGAACACCACTTCGCTGA-3’ |
| Mouse RPRM forward | 5’-TGCATTTCCAGCTGGGTCTT-3’ |
| Mouse RPRM reverse | 5’-GAGTCAAACGGTGTCACGGA-3’ |
| Mouse GAPDH forward | 5’-TGACCACAGTCCATGCCATC-3’ |
| Mouse GAPDH reverse | 5’-GACGGACACATTGGGGGTAG-3’ |
